# Halfway to self-sustainability: Reintroduced migratory European Northern Bald Ibises (*Geronticus eremita*) still need management interventions for population viability

**DOI:** 10.1101/2021.04.03.438331

**Authors:** Sinah Drenske, Viktoriia Radchuk, Cédric Scherer, Corinna Esterer, Ingo Kowarik, Johannes Fritz, Stephanie Kramer-Schadt

## Abstract

Northern Bald Ibis (NBI) have disappeared from Europe already in Middle Age. Since 2003 a migratory population is reintroduced in Central Europe. We conducted demographic analyses of survival and reproduction of 384 NBI over a period of 12 years (2008-2019). These data also formed the basis for a population viability analysis (PVA) simulating the possible future development of the NBI population in different scenarios. We tested life-stage specific survival rates for differences between these stages, raising types and colonies as well as the influence of stochastic events and NBI supplements on the population growth.

Stage specific survival rates ranged from 0.64 to 0.78. 61% of the mature females reproduce with a mean fecundity of 2.15 fledglings per nest. The complementary PVA indicated that the release population is close to self-sustainability with a given lambda 0.95 and 24% extinction probability within 50 years. Of the 326 future scenarios tested, 94 % reached the criteria of <5% extinction probability and population growth rates >1. In case of positive population growth, stochastic events had a limited effect. Of 820 sub-scenarios with different stochastic event frequencies and severities 87 % show population growth despite the occurrence of stochastic events.

Predictions can be made based on the results of the individual-based model as to whether and under what circumstances the reintroduced NBI population can survive. This study shows that a PVA can support reintroduction success that should work closely together with the project in the field for mutual benefit, to optimize future management decisions.

## Introduction

In conservation biology, species restoration plays an increasingly important role to counteract the currently ongoing dramatic decline of biodiversity, to assist the colonization of once widespread species and to restore whole populations that have become extinct (IUCN/SSC, 2013a; Destro et al., 2018; Pettorelli et al., 2018). Reintroductions are a key restoration method, defined by the IUCN as “the intentional movement and release of an organism inside its indigenous range from which it has disappeared”. Reintroductions commonly aim at minimizing interventions and allow for viable populations with lowest possible human support (Corlett, 2016).

Reintroduction efforts have been performed for an increasing range of species (Frey, 1992; Bennett et al., 2013; Fritz, Kramer, et al., 2017; Gray et al., 2017; Soorae, 2018), showing different levels of success. Impairments are mainly due to species interactions like predation, human-wildlife interactions and missing habitat suitability that all affect species’ viability and reproduction (Wimberger et al., 2009; Bennett et al., 2013; Robert et al., 2015). There is no general definition of reintroduction success (Robert et al., 2015), but it is recommended to calculate the population growth rate, population size and reproduction probabilities and the frequency of stochastic events, i. e. events that occur with a given probability and reduce the population size in the reintroduction site (Robert et al., 2015). Besides, it is of paramount importance to not only assess the long-term success of reintroductions under given constraints, but also to evaluate and rank the impact of management interventions taken in a retrospective way to be able to adjust management actions (Pereira & Navarro, 2015; Robert et al., 2015).

In this paper, we retrospectively analyse long-term demographic data (2008-2019) of a European migratory release population of the Northern Bald Ibis (*Geronticus eremita*, hereafter NBI) and assess the long-term success of the reintroduction alongside the impact of management measures in an individual-based model (IBM). In 2002, a European group of scientists, called *Waldrappteam* and headed by J. Fritz, started an NBI research project which in 2014 changed into an EU LIFE+ funded reintroduction project. Their mainly used translocation action is the Human-Led Migration. Chicks from zoo breeding colonies are raised by human foster parents at breeding sites north of the Alps and trained to follow a microlight airplane, which leads them to the wintering site (Fritz, Kramer, et al., 2017; see also Supplementary Material 1).

We hypothesize that at the given stage the NBI population can survive without further management and release. We predict that observed demographic rates will ensure population growth and do not differ between the colonies.

## Material & Methods

### Species and study area

The NBI is a threatened species, listed in the IUCN Red List as *critically endangered* from 1994 and down listed to *endangered* in 2018. The species is migratory, mainly insectivorous and reaches an age of up to 30 years in captivity. NBI are mainly seasonally monogamous and breed in colonies of up to hundreds of birds (Boehm et al., 2020). Juveniles learn the migration route by following conspecifics to the wintering ground, there they usually remain until they reach sexual maturity (Fritz, Unsoeld, et al., 2019). The last remaining wild population in the world changed to a sedentary lifestyle and lives year round at two breeding sites on the Atlantic coast of Morocco (Bowden et al., 2008).

The European release population consists of four breeding colonies in southern Germany (Burghausen, Überlingen) and Austria (Kuchl, Rosegg). The common wintering area is located in Italy, in the Tuscan nature reserve WWF Oasi Laguna di Orbetello (Fig. 1) (Fritz, Unsoeld, et al., 2019).

**Figure 1.**
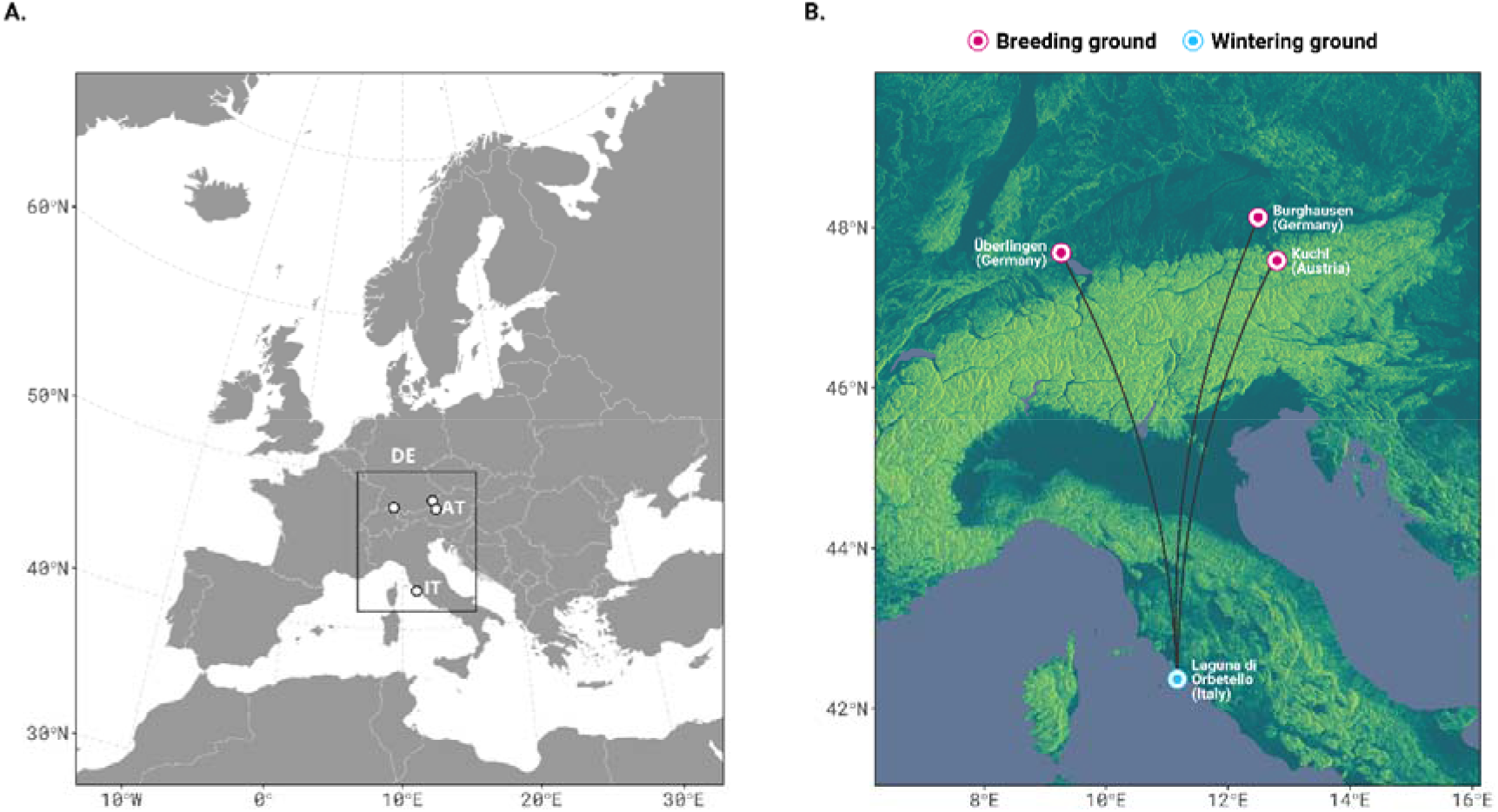
Study site of Northern Bald Ibis in Europe. a) Location of the study area in Europe. b) Closer look on two already established breeding sites (pink) in Burghausen (Bavaria, Germany) and Kuchl (State of Salzburg, Austria) and new colonies in Ueberlingen (Baden-Württemberg, Germany; pink) and Rosegg (Kärnten, Austria) with released NBI. The common wintering ground is the WWF Oasi Laguna di Orbetello in Tuscany, Italy (blue). Black lines: migration routes.

### Data collection & study design

384 NBIs which hatched between 2008 and 2019 are included in the study. Due to insensitive monitoring comprehensive data on genetically determined sex, raising type (either human-raised founder individuals FP, generation F0, or biological parent raised wild individuals BP, generations F1+), breeding colony and potentially date and cause of death or disappearance are available. Reproduction data include egg laying dates, clutch size, and pre-fledging survival probability. In the first years, spatio-temporal data are based exclusively on sight reports. From 2012 on, an increasing proportion of the birds carried GPS tags and from 2014 onwards, the whole population could be remotely monitored (Sperger et al., 2017).

Founder individuals are raised by human foster parents (FP). In autumn of their first year, they followed a microlight air-plane to the wintering ground where they were released (Fritz, Kramer, et al., 2017). Since 2011, chicks are raised in the wild by their biological bird parents (BP). They follow their conspecifics to the common wintering ground along the migration corridor, which was established during release of the founder individuals. At time of the study, some of the released founder individuals were not yet assigned to a breeding colony, as they have not yet migrated back to a breeding area.

The NBIs of all four breeding colonies are considered as one population, sharing the same wintering ground and separating for breeding, with genetic exchange (Fritz, Kramer, et al., 2017; Fritz, Wirtz, et al., 2017; Wirtz et al., 2018).

### Definition of demographic stages and demographic analyses

For the determination of demographic stages, the 384 NBI were assigned to four stages representing age classes with characteristic life-history events (Fig. 2; see also Supplementary Fig. 2). Stage 1 are juveniles from fledging until end of their first year of life; after the first autumn migration these birds usually remain at the wintering grounds. Stage 2 are juveniles in their second year of life; they usually stay year-round at the wintering grounds. Stage 3 are subadult birds in their third year of life; although not mature, most of them perform a partial or even full migration. Stage 4 are adults older than 3 years; these birds usually migrate and reproduce annually, even though some individuals may remain at the wintering ground.

**Figure 2.**
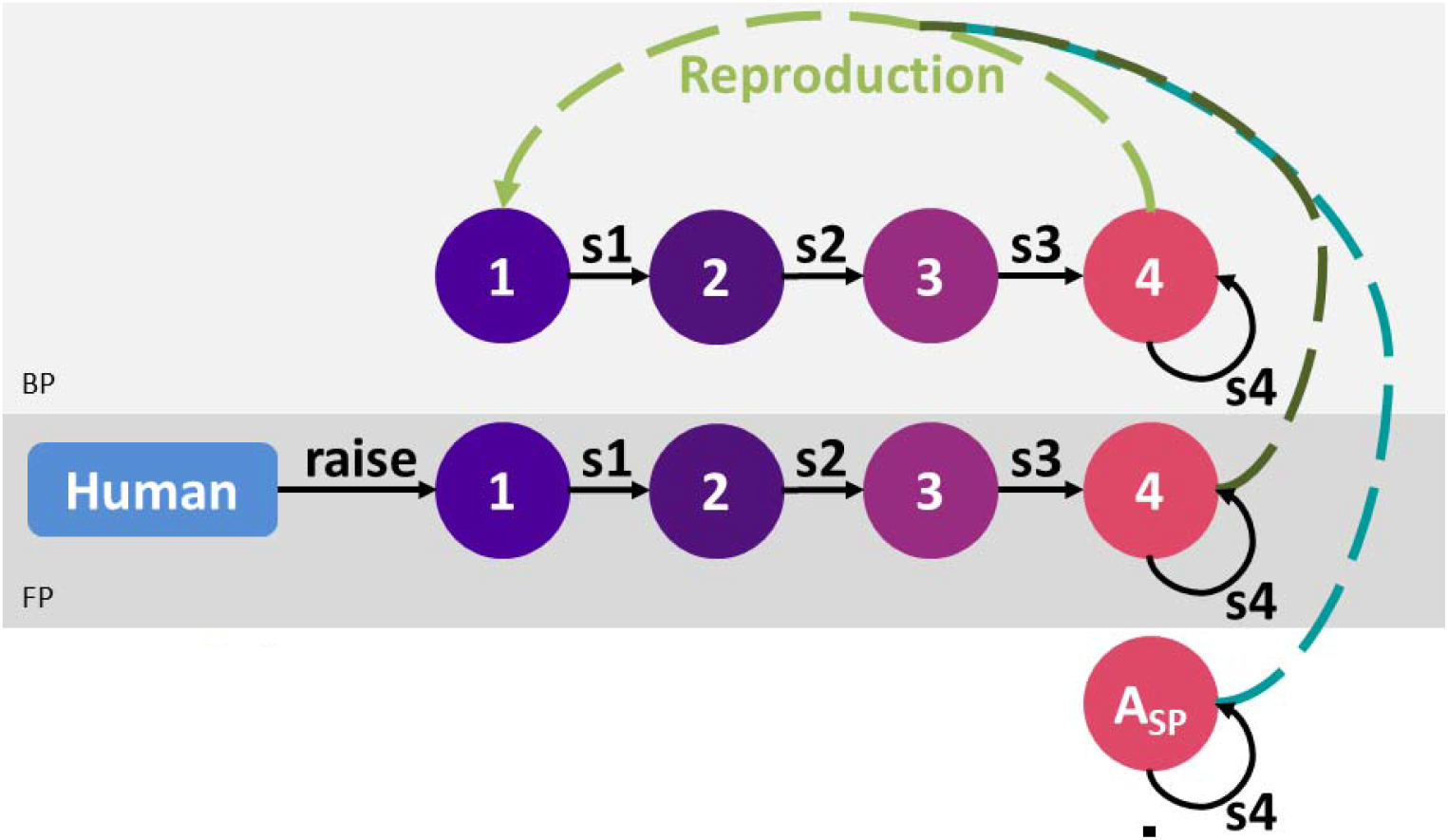
Life cycle graph of the Northern Bald Ibis. Life stages are in circles, for definition see text; s1 to s4 are the survival probability of the regarding stages; light grey section: biological parent (BP) raised proportion of the population; dark grey section: human foster parent (FP) raised part of the population; A_SP_: temporarily added females. Only the female part of the population is considered for the simulations.

For the calculation of survival probabilities per stage, sex, colony and raising type we used the Kaplan-Meier estimator (Kaplan & Meier, 1958) with the ‘survival’-package v. 3.2-7 in R v. 4.0.3 (Therneau, 2015; R Core Team, 2020). We estimated survival probabilities for fledglings around 45 days of age. Adult birds that were only temporarily added during the breeding season to improve breeding success were discarded from the calculations of the survival probabilities. We used a likelihood ratio test (LRT) to test for significance between different classes in Cox proportional hazard models (Kleinbaum & Klein, 2012). We also calculated hatching rate and fledging rate.

Fecundity was calculated as the mean number of fledglings per nest. At the start-up of new founded colonies, adult males and females were temporarily added at the breeding site to compensate for missing mates and to enable the birds to reproduce. The rate of temporarily added adults was successively reduced and completely terminated at the end of the data period.

### Population Viability Analysis

For the Population Viability Analysis (PVA) we focus our analyses and population projections on the female half of the population (underlying a sex-ratio of 1:1). We tested how different values of demographic rates used for the four stages, as analysed from field data, influence population trajectories and population viability.

We analysed the population trajectories per scenario (see below) and calculated the extinction probability as the number of runs of 100 repetitions per scenario where the population went extinct (0 individuals) within the simulated 50 years (*P_EXT_50_*), and the intrinsic growth rate of the population (lambda), calculated as the mean of the annual finite rate of change in population size. Then, we analysed the distribution of the input parameters of demographic rates in scenarios where lambda > 1 and *P_EXT_50_* ≤ 5%. In addition, we ran generalized linear models (GLMs) and conducted an analysis of variance (anova) to rank the contribution of demographic rates in the different stages on lambda as response variable, using a gamma error structure with inverse link. Besides, we calculated how often each combination of stochastic event frequency (5-20%) and severity (5-25%) and each combination of numbers of supplements (15 or 30) and time span of intervention (4 or 7 years) occurred in sub-scenarios resulting in lambda > 1 and *P_EXT_50_* 5%.

The model documentation follows the TRACE documentation framework (Grimm et al., 2014; Supplementary Material 3). The model description follows the ODD protocol (Grimm et al., 2006, 2010). An individual-based model (IBM) was implemented in NetLogo 6.0.3 (Wilensky, 1999). The corresponding NetLogo and R scripts are provided online (Supplementary Material 6).

### Reproductive Rates

For the PVA we only considered female fledglings. From fledglings with unknown sex 50% were included. For the analysis we included all adult females in the release population (stage 4), irrespective whether they were breeding or not, plus adult females which were temporarily added at the breeding site to compensate for missing mates. Thus, different to RRNest this reproductive rate is an expression of breeding probability, it indicates how likely it is that a female will breed.

Three different reproductive rate (RR) were included into the PVA, depending on the fledglings that were included: RR_Baseline_ includes fledglings raised by adult wild females (stage 4); RR_Status quo_ includes in addition female fledglings raised by temporarily added females; RR_All chicks_ includes in addition female fledglings raised by human foster-parents.

RR_Baseline_ was also calculated separately for the two colonies Burghausen and Kuchl (RR_B_ and RR_K_) and separately for females raised by bird parents (RR_BP_) and human foster-parents (RR_FP_). We also implemented GLMMs (generalized linear mixed models) using a Poisson error structure and maximum likelihood fit, to investigate if differences in reproduction are significant.

### Scenario development

We analysed the model for two different types of scenarios:

(I) The management scenarios (MScen) compare situations where no management took place (baseline scenario S_Base_), i.e. release of fledglings or temporary release of adults, to scenarios where baseline demographic rates were improved by 10%, 25% and 100%. We combined three levels of survival rates at each stage and four levels of RR in a full factorial design. Additionally, we combined RR_status quo_ and RR_all chicks_ with baseline survival probabilities, which resulted in 326 MScen (Table 1, Supplementary Material 4). Only for the simulation of S_Base_ we draw the demographic rates from the probability distributions defined by mean and standard deviation to test Hypothesis 1 (H1). For all other scenarios we used the calculated mean values of demographic rates. Besides, we chose 14 scenarios of special interest for closer examination (Table 1). All parameter combinations were within the scope of realistic management measures to improve survival and reproduction.

**Table 1.** In the upper part, the table shows the given empirical values for survival per stage and for the three reproductive rates as defined in the text. These values and their improvements by 10%, 25% or 100% were used in different combinations for the NetLogo simulation of 14 management improvement scenarios, as outlined in the lower part of the table. Improvement of the empirical values are indicated by bold numbers. The resulting Lambda and extinction probability (*P_EXT_50_*) are shown in the two right columns. Please note that reproductive rate comprises not only the reproducing females but all adult females in stage 4 and takes into account only to female offspring. In brackets: Standard deviation.

| Results and PVA scenarios/stage | s1 | s2 | s3 | s4 | Reproductive rate (females) | Lambda | P<sub>EXT_50</sub> |
| --- | --- | --- | --- | --- | --- | --- | --- |
| Empirical values |  |  |  |  |  |  |  |
| RRBaseline | 0.64 (±0.36) | 0.74 (±0.35) | 0.69 (±0.35) | 0.78 (±0.14) | 0.53 (±0.17) |  |  |
| RRstatus quo |  |  |  |  | 1.41 (±0.81) |  |  |
| RRAll chicks |  |  |  |  | 3.97 (±2.66) |  |  |
| Management improvement scenarios I - 14 scenarios of special interest |  |  |  |  |  |  |  |
| Baseline (RRBaseline) | 0.64 | 0.74 | 0.69 | 0.78 | 0.53 | 0.95 (±0.030) | 24% * |
| +10% RR | 0.64 | 0.74 | 0.69 | 0.78 | **0.58** | 0.97 (±0.015) | 3% |
| +25% RR | 0.64 | 0.74 | 0.69 | 0.78 | **0.66** | 0.99 (±0.013) | 0% |
| +100% RR | 0.64 | 0.74 | 0.69 | 0.78 | **01.Jun** | 1.07 (±0.005) | 0% |
| +10% s1 | **0.70** | 0.74 | 0.69 | 0.78 | 0.53 | 0.97 (±0.020) | 1% |
| +25% s1 | **0.80** | 0.74 | 0.69 | 0.78 | 0.53 | 0.99 (±0.014) | 1% |
| +10% s4 | 0.64 | 0.74 | 0.69 | **0.86** | 0.53 | 1.02 (±0.007) | 0% |
| +25% s4 | 0.64 | 0.74 | 0.69 | **0.98** | 0.53 | 1.10 (±0.004) | 0% |
| +10% s1-s4 | **0.70** | **0.81** | **0.76** | **0.86** | 0.53 | 1.05 (±0.006) | 0% |
| +25% s1-s4 | **0.80** | **0.92** | **0.86** | **0.98** | 0.53 | 1.19 (±0.005) | 0% |
| +10% s1-s4 and RR | **0.70** | **0.81** | **0.76** | **0.86** | **0.58** | 1.07 (±0.005) | 0% |
| +25% s1-s4 and RR | **0.80** | **0.92** | **0.86** | **0.98** | **0.66** | 1.23 (±0.004) | 0% |
| Status quo (RRstatus quo) | 0.64 | 0.74 | 0.69 | 0.78 | Jän.41 | 1.13 (±0.006) | 0% |
| All chicks (RRAll chicks) | 0.64 | 0.74 | 0.69 | 0.78 | Mär.97 | 1.40 (±0.012) | 0% |

\* For a simulation across 100 years the extinction probability would be 87%.

(II) The stochastic event and juvenile supplement scenarios (SJS) assess reproduction improvement by the supplemental release of FP juveniles (F0). For these SJS we crossed S_Base_ and the MScen of special interest where lambda >1 and *P_EXT_50_* ≤ 5% (9 MScen, see results) with 4 levels of stochastic event frequency (5, 10, 15, 20), 5 levels of severity (5, 10, 15, 20, 25; additional mortality per stage), 2 levels of the number of supplements (15, 30) and 2 levels of the time for supplementing individuals (4 or 7 years).

## Results

### Demography

From the 384 individuals, 195 were males and 184 females, what gives a sex rate of 1.00:0.94 (m:f). For 5 juveniles, sex was not determined at time of data analysis. Concerning the raising type, N=162 birds were raised in the wild by bird-parents (42%; BP), N=213 individuals by foster-parents (55%; FP) and N=9 supplemented as juveniles (2%). Concerning the breeding colonies, N=127 individuals belonged to Burghausen (33%), N=89 to Kuchl (23%), N=89 to Überlingen (23%), N=22 to Rosegg (6%) and N=57 juveniles were not yet assigned at time of the data collection. 232 birds died during the data collection period; out of that N=123 during stage 1 (53%), N=45 during stage 2 (19%), N=33 during stage 3 (14%) and N=31 during stage 4 (13%; see also Supplementary Fig. 1).

At the end of 2019, 152 individuals were alive including 9 individuals which were delivered to zoos for various reasons. Thus, the release colony at end of the data collection consisted of 143 individuals, 69 males and 74 females (sex rate of 1.00:1.07). Concerning the life stage, N=71 individuals belonged to stage 1 (50%; 37f, 34m), N=28 to stage 2 (20%; 11f, 17m), N=13 to stage 3 (9%; 8f, 5m) and N=31 to stage 4 (22%; 18f, 13m). Concerning the colonies, N=33 bird belonged to the colony Burghausen (23%), N=33 to Kuchl (23%), N=51 to Überlingen (36%) and N=15 to Rosegg (10%); 11 individuals (8%) were not yet allocated to a colony. Concerning the raising types, N=68 birds (48%) were BP raised, N=73 FP raised (51%) and N=2 birds (1%) were supplemented.

#### Survival

Survival was calculated for all individuals from fledging onwards. According to the cox proportional hazard model survival rates between stages across the whole population did not differ significantly (LRT, N = 736, 3.05, *p* = 0.4). Neither did they differ significantly between the sexes (LRT, N = 379, 0.8, *p* = 0.4). According to the Kaplan-Meier estimator, survival probability was for stage 1 0.64 (± 0.36) for stage 2 0.74 (± 0.35), for stage 3 0.69 (± 0.35) and for stage 4 0.78 (± 0.14). The cumulative survival probability until sexual maturity (end of stage 3) was 0.33 (± 0.23). Concerning pre-fledging survival, 72% of the eggs hatched and 83% of the hatched chicks fledged (Table 2).

**Table 2.** Breeding statistics of the *Waldrappteam* population: in brackets: Standard deviation.

| Year | Number of<br>nests per<br>year | Developmental stage |  |  |  |  |  |
| --- | --- | --- | --- | --- | --- | --- | --- |
|  |  | Eggs |  | Hatchlings |  | Fledglings |  |
|  |  | Total<br>number | Mean per nest | Total number | Mean per nest | Total<br>number | Mean per nest |
| 2011 | 1 | 3 | 3.00 (NA ) | 3 | 3.00 (NA ) | 3 | 3.00 (NA ) |
| 2012 | 5 | 11 | 2.20 (± 1.30) | 9 | 1.80 (± 1.10) | 8 | 1.60 (± 1.34) |
| 2013 | 7 | 23 | 3.29 (± 0.76) | 9 | 1.29 (± 1.25) | 6 | 0.86 (± 1.21) |
| 2014 | 8 | 23 | 2.88 (± 0.64) | 15 | 1.88 (± 0.64) | 13 | 1.63 (± 0.74) |
| 2015 | 6 | 19 | 3.17 (± 0.75) | 19 | 3.17 (± 0.75) | 17 | 2.83 (± 0.75) |
| 2016 | 6 | 30 | 5.00 (± 2.45) | 15 | 2.50 (± 1.64) | 13 | 2.17 (± 1.47) |
| 2017 | 9 | 35 | 3.89 (± 1.62) | 24 | 2.67 (± 1.41) | 18 | 2.00 (± 1.66) |
| 2018 | 10 | 37 | 3.70 (± 0.48) | 32 | 3.20 (± 0.79) | 26 | 2.60 (± 0.52) |
| 2019 | 14 | 53 | 3.79 (± 1.12) | 43 | 3.07 (± 1.64) | 37 | 2.64 (± 1.50) |
| **Total** | **66** | **234** | **3.43 (± 0.79)** | **169** | **2.51 (± 0.70)** | **141** | **2.15 (± 0.70)** |
| Survival |  |  |  | 0.72<sup>1</sup> |  | 0.83<sup>2</sup> |  |

<sup>1</sup> percent of eggs hatched; <sup>2</sup> percent of hatchlings fledged.

Survival between the two raising types FP and BP differed (LRT, N = 375, 8.26, *p* = 0.004; Figure 3), also per stage and per raising type (LRT, N = 715, 10.17, *p* = 0.04). In stage 1 FP raised individuals showed a higher survival rate (FP = 0.73 ± 0.37, BP = 0.52 ± 0.30), while in stage 4 BP raised adults showed a higher survival rate (FP = 0.72 ± 0.22, BP = 0.92 ± 0.17; Table 1). The survival rates in the two colonies Burghausen and Kuchl did not significantly differ, neither for the whole colonies (LRT, N = 216, 1.56, *p* = 0.2) nor per stage and per colony, respectively (LRT, N = 453, 5.36, *p* = 0.3; Table 1).

**Figure 3.**
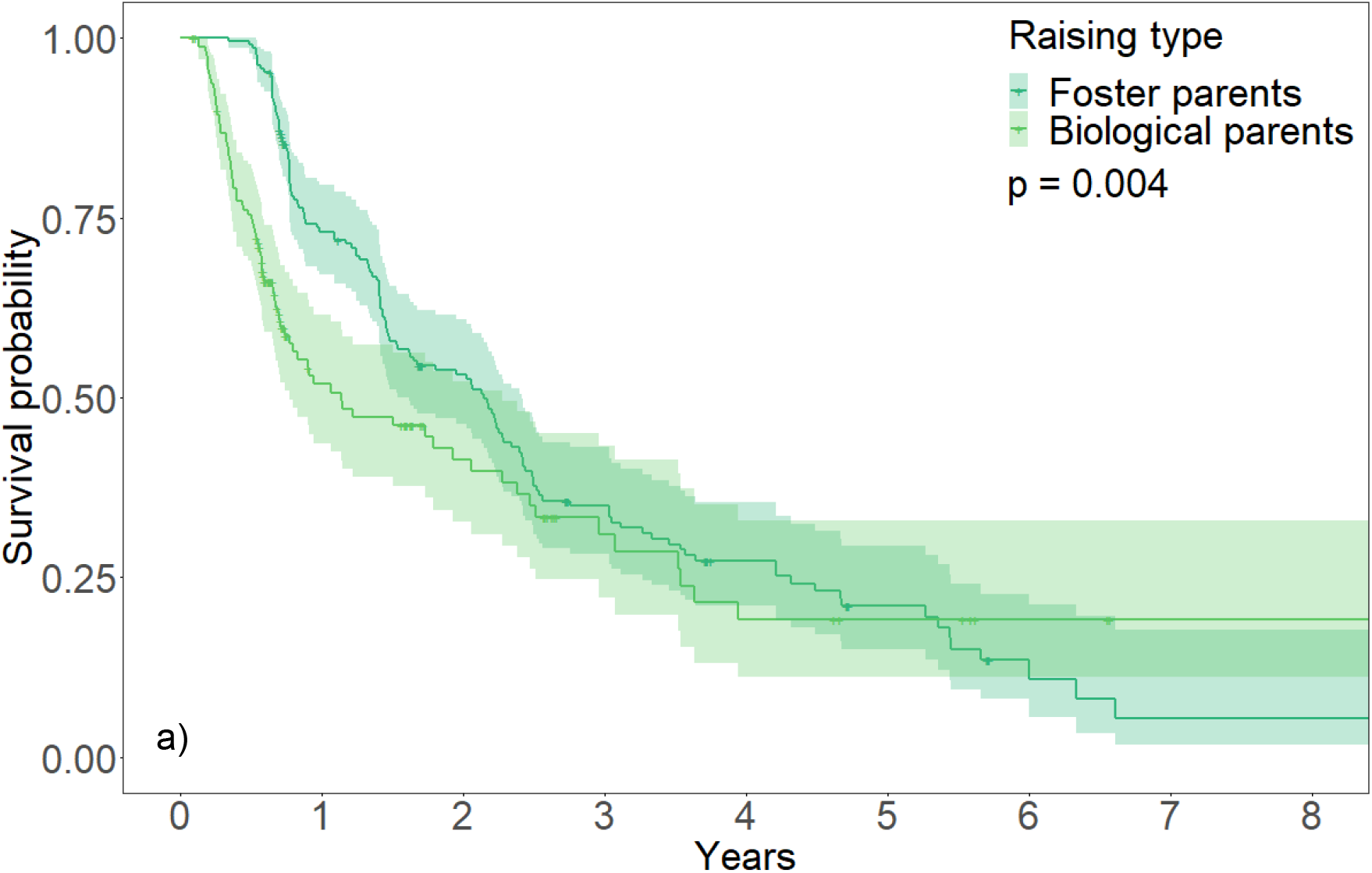
Survival plot of the raising types. The p-values are the results of the LRT. Shades of the lines represent the standard deviation.

#### Fecundity

The number of nests increased steadily with 1 nest in 2011 and 14 nests in 2019, with a mean of 3.43 (± 0.79) eggs per nest (Table 2). Overall fecundity was 2.15 fledglings per nest (± 0.70). Fecundity rates of females raised by human-foster parents and females raised by their bird parents (BP) in the wild did not differ significantly (z = 0.68, p = 0.50), neither fecundity rates among the colonies Burghausen and Kuchl (z = 0.60, p = 0.55).

The annual rate of reproducing adult females (s4) was 0.61 (± 0.20). From a total of 27 potentially reproductive females N=13 (48%) never reproduced till end of the data collection period, N=4 (15%) reproduced once, N=4 (15%) twice, N=5 (19%) three times and N=1 (4%) 6 times.

### Population Viability Analysis

#### Reproductive Rate

Over the whole period of data collection, a total of 62 individuals reached stage 4. Analysing 31 females of these 62 individuals of both sexes in stage 4 as potential mothers resulted in a RR_Baseline_ of 0.53 (± 0.17; Table 1) (wild BP and FP mothers and their female fledglings), RR_Status Quo_ of 1.41 (± 0.81) (additionally female fledglings from temporarily added females included) and RR_All Chicks_ of 3.97 (± 2.66) (additionally human raised and released female fledglings included). These reproductive rates were used for the PVA.

#### Management scenarios (MScen)

Without further management and translocation measures, as assumed for the baseline scenario, the PVA indicates a lambda of 0.95 (± 0.030), below the 1.0 threshold with 24% extinction probability within 50 years (Table 1).

The full factorial design resulted in 326 MScen (Supplementary Material 4). Across all 326 MScen lambda ranged between 0.95 and 1.40. Lambda was >1 and *P_EXT_50_* ≤ 5% in 308 out of 326 MScen (94 %), in all these 308 MScen the population did not go extinct. In a first analysis we investigated the frequencies of MScen with different demographic rates out of 306 of 308 MScen with positive population development, to understand how demographic rates affect population viability (Fig. 4). The two MScen “Status quo” and “All chicks” were not considered here, because they were only simulated once with baseline survival values and the respective RR. In the 306 MScen with positive population development only s4 shifted to increasing survival values, as given by the improved MScen. An increase of s4 by 10% or 25% increased the viability disproportionately (each 108 times) compared to the baseline value (90 times). These results indicate the importance of survival of the reproductive stage for population viability. This is supported by the GLMs and the ANOVA ranking the effect of the survival rates (deviance d of ANOVA: d_s1_ = 0.05, d_s2_ = 0.05, d_s3_ = 0.05, d_s4_ = 0.68). Here, s4 had the largest effect, together with reproductive rates (Supplementary Material 5).

**Figure 4.**
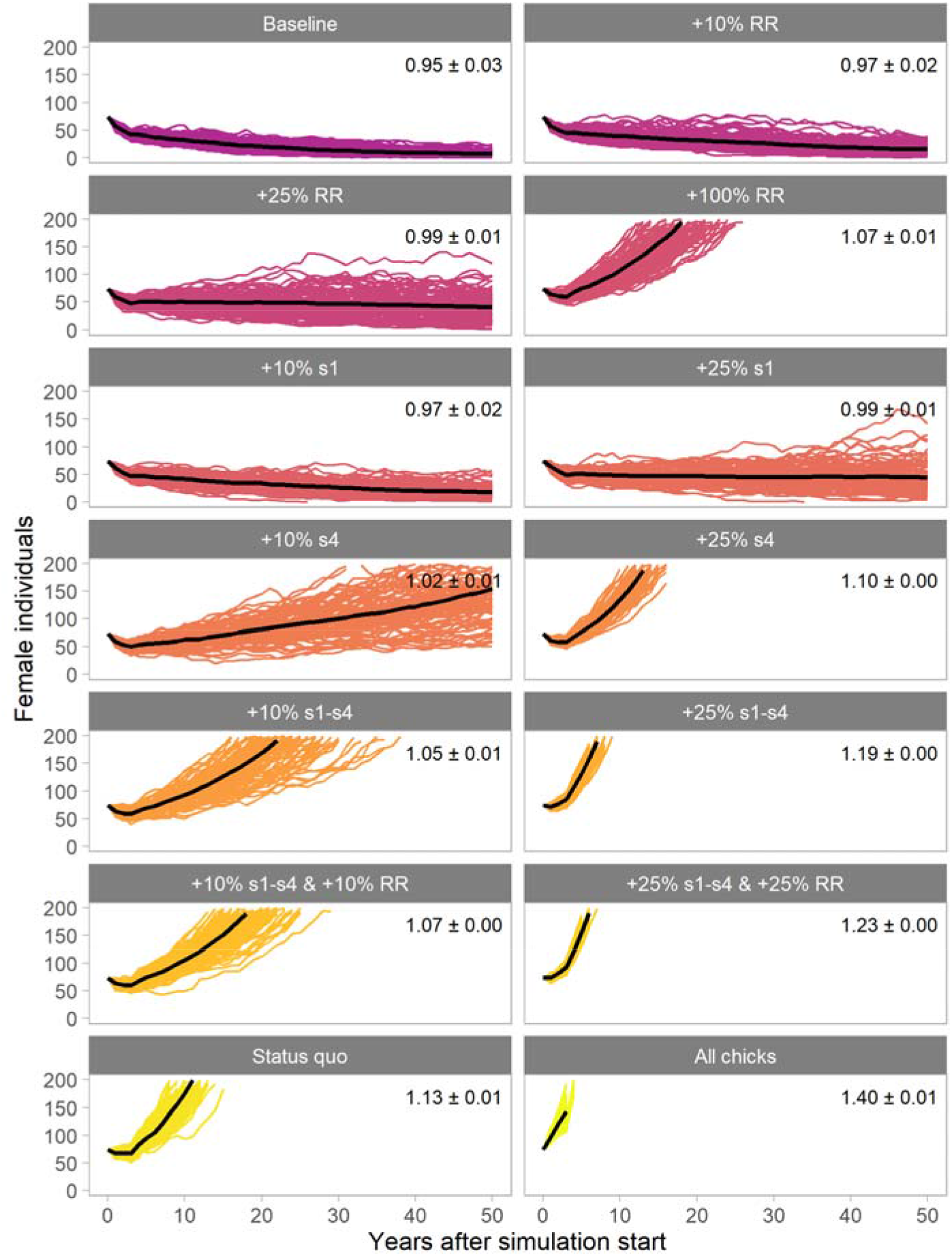
Number of females per year in the 14 scenarios of special interest. Each line of a facet corresponds to a run of the respective scenario. The bold black line represents the average of all 100 runs per scenario. Lambda ± standard deviation are specified for each scenario. The description of each scenario can be found in the heading of the facet. s1-s4 are the survival probabilities, RR is the reproductive rate. The starting point is 74 females.

All MScen where RR was increased by 100% showed positive population development (81 times). Only one MScen where RR was increased by 25% was rejected, i. e. the one where no survival rate was increased (i. e. baseline values for s1-s4, RR +25% MScen; Table 1). Still, 88% of the MScen with RR_Baseline_ showed positive population development (71 times). The reproductive rate had a strong effect on lambda, too (d_*RR*_ = 0.58) in the GLMs.

In the second analysis 14 MScen of special management interest were analysed (Table 1, Fig. 5). For nine out of these, lambda was >1 and *P_EXT_50_* ≤ 5%. If only RR was increased, positive population growth only occurred from an increase of at least 100% (“+100% RR”, “Status quo”, “All Chicks”), i. e. a minimum of one female fledgling per female. Increasing only the survival of juveniles s1 did not lead to positive population growth but increasing adult survival s4 did. An increase in all survival values s1-s4 also led to positive population growth; these effects were reinforced by an additional increase in the reproductive rate.

**Figure 5.**
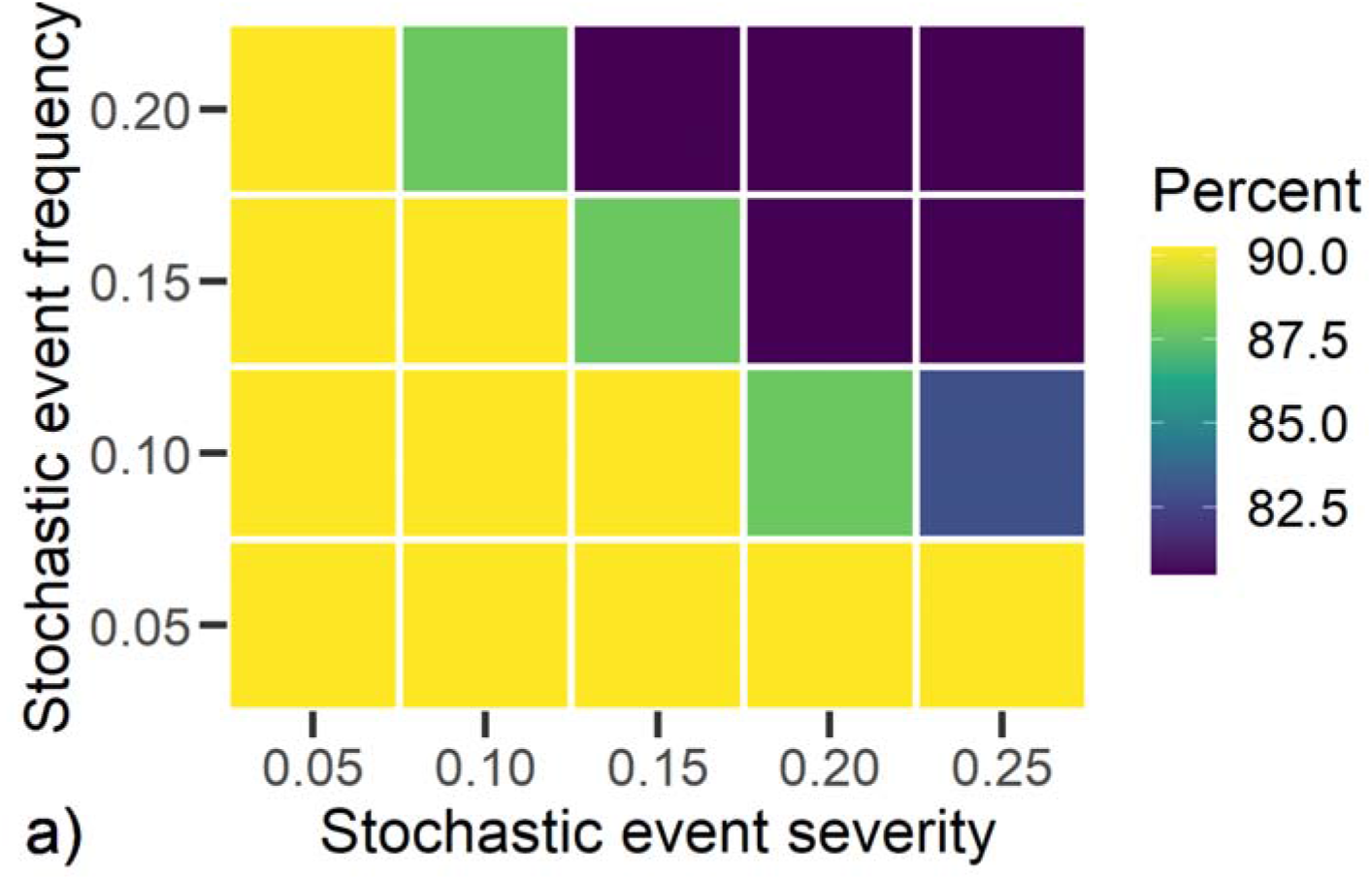
Percentage of the respective combination of stochastic event frequency and severity in the 714 sub-scenarios where lambda >1 and extinction probability *P_EXT_50_* ≤ 5%. In all sub-scenarios, each combination occurred 41 times.

#### Stochastic event and juvenile supplement scenarios (SJS)

Here we developed 820 scenarios, taking 11 MScens: nine MScen of special interest where lambda >1 and *P_EXT_50_* ≤ 5%, the SBase scenario and once the “All chicks” scenario without the supplements, as the supplements are already included in the RR_All Chicks_ (see under Supplementary Material 2). The number of SJS is given by the equation in Supplementary Material 4. Lambda was between 0.91 and 1.40. In 714 out of 820 SJS (87 %) lambda was >1 and *P_EXT_50_* ≤ 5%. Baseline survival values led to positive population growth in 74-79% of the SJS. Increase of survival of adults (stage 4) by 10% led to positive population growth in 89% of the SJS, while 10% survival increase in other age categories led to population growth in 100% of the SJS considered. Increasing baseline survival by 25% led to positive population growth in 100% of the SJS.

Each combination of stochastic event frequencies and severities occurred 41 times in all 820 SJS. 90% of the SJS with low frequency (5%) and severity (5%) of stochastic events and also 80% of the SJS with high frequency (20%) and severity (25%) of stochastic event resulted in positive population development (Fig. 5). 88% of the SJS with 30 supplemented juveniles per year over 7 years and also 86% of the SJS with 15 supplements per year over 4 years resulted in positive population development.

## Discussion

This paper presents an analysis of demographic development, survival, and reproduction of a reintroduced Northern Bald Ibis population. It covers a period of 12 years, including the late phase of a research project (2008-2013), where small number of birds were released, and the following six-year period of the LIFE+ funded reintroduction project (2014-2019).

The comprehensive 12-year data set allowed a Population Viability Analysis (PVA) of the future population development under different scenarios. PVA is a powerful mean to evaluate the effectiveness of reintroduction and management scenarios. Such models based on the best possible evidence are recommended in the IUCN reintroduction guidelines to decide on further management (IUCN/SSC, 2013b).

### Survival

Survival rate of the *Waldrappteam* population is ranging from 64% for juveniles (stage 1) to 78% for adults (stage 4). A matching adult survival rate of 81% is published for a small migratory NBI relict population, which was discovered in the Middle East in 2002 (Serra et al., 2014). Since the Moroccan birds are not marked, their survival rate can only be estimated due to the annual counts of the pre-breeding population and the number of fledglings (Bowden et al., 2008). This results in an overall annual survival rate of 77%, what is also in accordance with the calculated rates of the *Waldrappteam* population.

The cumulative survival rate of the *Waldrappteam* population until sexual maturity (stage 1 to 3) was 33%. In the Middle East relict population, the count of departing fledglings and arriving new breeders at the breeding site in Syria indicates 14% survival rate until sexual maturity (Boehm et al., 2020), what it less than half the rate of the *Waldrappteam* population. Satellite-tracking of the Middle East juveniles revealed, that they did not arrive at the common wintering site in Ethiopia, because they lost the company of experienced conspecifics and died at the Arabic Peninsula (Serra et al., 2014). This low survival rate of juveniles was a major cause for the extinction of this population in 2013, despite international conservation and translocation efforts (Fritz & Riedler, 2010; Bowden et al., 2012; Serra, 2015).

In our population, first-year survival of foster-parent raised and released juveniles (73%; FP) was significantly higher compared to bird-parent raised juveniles (52%; BP). This contrasts the widespread experience that released individuals have comparably low survival rates, particularly in the first period after release, due to translocation and competition induced stress, dispersal, diseases or predation (Parker et al., 2013), wherein this is true in particular for captive-bred individuals, probably because they lack individual experiences (Mathews et al., 2005). Our birds undergo an extensive pre-release training, which provides them with essential experiences regarding navigation, flight techniques, weather conditions, aerodynamics and predator avoidance (Portugal et al., 2014; Voelkl & Fritz, 2017; Fritz, Unsoeld, et al., 2019). This is assumed to improve post-release viability (Alonso et al., 2011; Houser et al., 2011; Zhang et al., 2017). In a Spanish NBI reintroduction project (*Proyecto Eremita*), the mean first-year survival rate for released juveniles was substantially lower with 31% (Boehm et al., 2020; data from 2004-2018), compared to the *Waldrappteam* population. We assume that minor pre-release training and unguided autumn dispersal (Muñoz & Ramírez, 2017) account for the strongly deviating survival rate of the juveniles released in Spain (Fritz, Unsoeld, et al., 2019).

The survival rate from hatching to fledging in the *Waldrappteam* population is 83%, what is substantially higher compared to a given value of 47% for the Morocco wild population (Bowden et al., 2003). We assume this difference to be mainly related to the quality of the feeding habitats. For the European birds, studies indicate a high feeding efficiency and a correspondingly high abundancy of food animals (mainly worms and larvae) in the soil of meadows and pastures as the preferred feeding habitats (Zoufal et al., 2007; Fritz et al., 2017) while the feeding habitats at the Atlantic coast in Morocco consist of semi-natural steppes with sparse cover of perennial and annual vegetation and low availability of fresh water sources (Bowden et al., 2008). Accordingly, position data indicate a small activity range of about two kilometres radius around the nesting sites for the European birds during the breeding season (Fritz et al., 2016) while the foraging area of the two Moroccan colonies covers a strip of about 4 km inland along 50 km coastline. The fledgling survival rate in Morocco was even lower in the 90’s before supplementary fresh water was provided near to the breeding grounds from 1998 on (Smith et al., 2008).

### Fecundity

In the *Waldrappteam* population 61% of the mature females (stage 4) actually reproduce. The majority of the non-reproducing females remain at the wintering site. In comparison, 63% of the *full-grown* birds in the Moroccan population reproduce (Bowden et al., 2008). These rates are not directly comparable, because the Moroccan data are based on counts of un-marked birds. But the data indicate that also in this sedentary population with all birds on site a similar proportion of the adults is not reproducing.

In the *Waldrappteam* population, the average annual fecundity was 2.15 fledged chicks per nest. Table 3 outlines fecundity values for other NBI populations, based on data published in (Boehm et al., 2020). The values vary considerably, with 1.23 fledged chicks per nest in the wild sedentary Moroccan population and 0.97 in the Spanish *Proyecto Eremita* population (release sedentary). The highest rate with 2.24 refers to the semi-wild managed population at Zoo Rosegg in Austria. The comparison indicates that fecundity in the *Waldrappteam* population is at the upper end, which in consistence with the high survival rate from hatching to fledging (83%) and indicates a high quality of the breeding habitats. The rate of 1.67 for a Chinese sedentary release population of the Crested ibis (*Nipponia nippon*) lies in the range of the NBI populations.

**Table 3.**
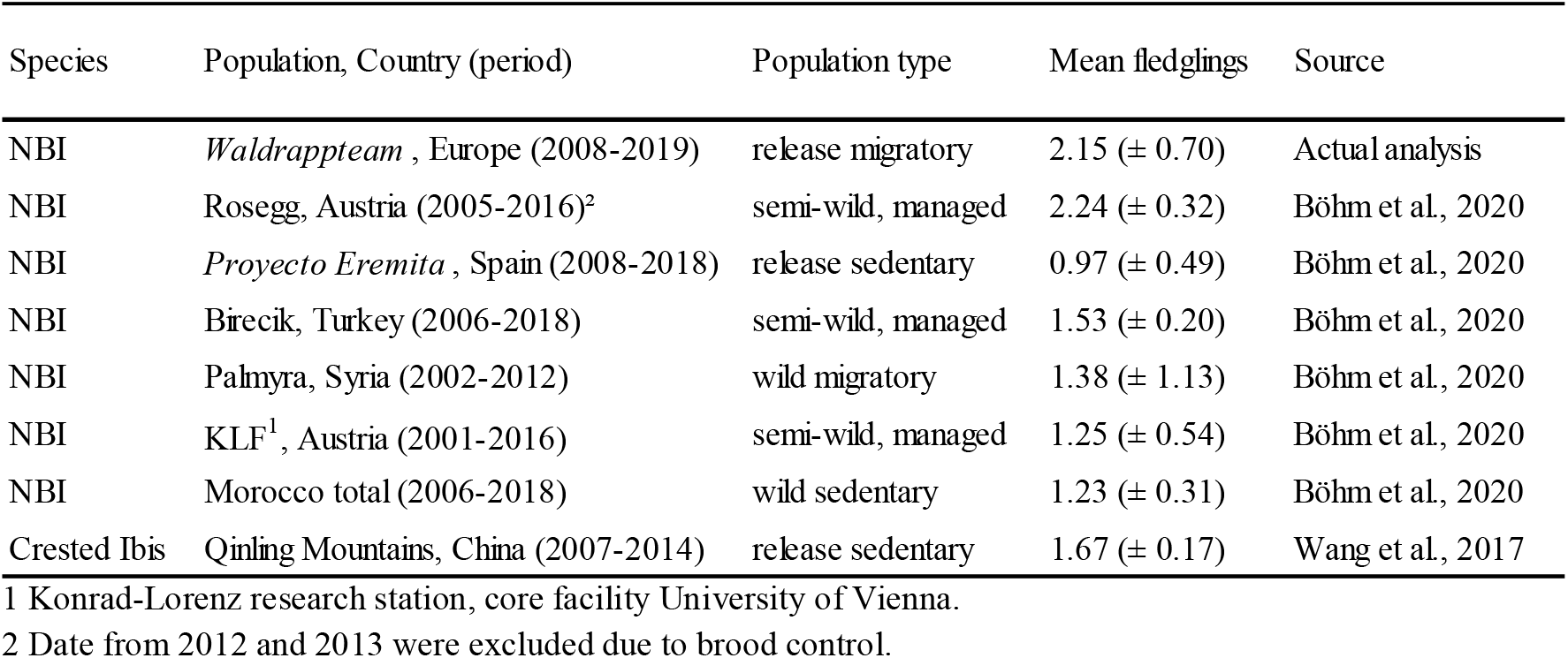
Comparative statistics on fecundity with mean number of fledglings per nest. SD = standard deviation. Bottom line Crested Ibis (*Nipponia nippon*).

### Population Viability Analysis

Due to the outcome of the PVA baseline scenario, with lambda 0.95 and 24% extinction probability within 50 years, the hypothesis that this reintroduced NBI population can survive without further management and release at the end of the data collection period (2019) is to be rejected. This result is not unexpected because the reintroduction project started just in 2014 and is still ongoing. Nonetheless, the population is already close to transition into a state of self-growth (with lambda >1.0) what also coincides with the comparatively good survival and fecundity values.

The baseline reproductive rate relates the actual number of fledged female offspring to the entirety of adult females (s4) in the population, including the 39% non-reproducing females. It is an expression of the reproductive potential in the population. The given baseline rate of 0.53 can be significantly increased by translocations measures, up to a rate of 3.97 (RR_All_ Chicks). As indicated by the management improvement scenarios continuing with the given translocation measures has the most significant immediate effect on lambda and extinction probability (Table 1). In comparison, an increase in the survival rate has a relatively small effect on lambda, where improvement in the survival rate of adults (s4) has the greatest impact. This is common knowledge and has also been confirmed for other long-lived species (Lebreton & Clobert, 1991; Lampila et al., 2006; Pistorius et al., 2006; Schaub et al., 2009; O’Shea et al., 2011). Sustainable improvement of the survival on long-term should be an objective for the project, even though intraspecific comparison indicates that in the *Waldrappteam* population survival values are already in a good range for all age categories.

The simulation of stochastic event scenarios with different frequency and severity led to only a slight impairment of the population development. The simulations also indicated that increasing survival or fecundity could mitigate the effect of stochastic events.

## Conclusion

Despite comparatively good survival and fecundity data the PVA indicated that the *Waldrappteam* population needs further management and translocation. It is important at this stage to plan the kind and duration of further management and translocation measures based on a quantitative, systematic analysis.

According to the modelling outcome, a major focus of future management of the release population will be on further improvement of the survival rates with particular focus on adults (stage 4). This will mainly be achieved by the implementation of measures against the major mortality causes, illegal hunting in Italy and electrocution on unsecured power poles at the breeding sites (Fritz, 2015; Fritz et al., 2019). Also, the proportion of reproducing females (61%) should be increased by the establishment of a breeding site south of the Alps which can be easily reached in spring without crossing the Alpine chain.

An interspecific comparison of the demographic data did not indicate adverse effects of the migratory lifestyle in the *Waldrappteam* population. The survival rate of adults is similar to sedentary populations and the survival rate of released juveniles is even substantially higher in the migratory population (73%) compared to a sedentary population in Spain (31%).

The high fecundity rate in the *Waldrappteam* population (2.15) is presumably a consequence of the migratory behaviour that allows the adults to exploit rich northern feeding grounds during the reproduction period.

The actual *International Single Species Action Plan* for the NBI (Bowden 2015) does not include modelling, neither as an objective nor as a recommendation. This indicates that systematic, quantitative approaches, such as the PVA, are still not well established and recognizes as a strategic tool for animal conservation. The study presented here show that the PVA could essentially contribute to the improvement of conservation measures for this and other endangered animal species and populations.

The model could be improved by including movement of the NBI in the PVA assessment to build a spatially explicit population simulation model accounting for differences in the migration routes from and to the colonies. In addition, a thorough assessment of the factors significantly influencing the viability of the NBI in the colonies would be an asset, such as the availability of food or nesting sites, the weather in the breeding areas, during the migration and in the winter area (Schaub et al., 2005).

## Supporting information

Supplementary Material 3 TRACE

## Author contribution

SD, SKS and JF conceived and designed the study, SD and VR analysed data, SD led the writing, CS supported programming, and CE collected field data. SKS, JF, IK and VR substantially contributed to the writing.

## Acknowledgements

The Waldrappteam is co-financed with a 50% contribution from the LIFE financial instrument of the European Union (LIFE+12-BIO_AT_000143, LIFE Northern Bald Ibis). SD, VR, CD and SKS are associated with the DFG funded research training group, BioMove□ (RTG 2118-1).

## Ethical standards

This research followed the *Oryx* guidelines on ethical standards.

## References

Alonso, R., Orejas, P., Lopes, F. & Sanz, C. (2011) Pre-release training of juvenile little owls Athene noctua to avoid predation. Animal Biodiversity and Conservation, 34, 389–393.

Bennett, V.A., Doerr, V.A.J., Doerr, E.D., Manning, A.D., Lindenmayer, D.B. & Yoon, H.J. (2013) Causes of reintroduction failure of the brown treecreeper: Implications for ecosystem restoration. Austral Ecology.

Boehm, C., Bowden, C.G.R., Seddon, P.J., Hatipoġlu, T., Oubrou, W., El Bekkay, M., et al. (2020) The northern bald ibis Geronticus eremita: history, current status and future perspectives. Oryx, 1–13.

Bowden, C.G.R., Aghnaj, A., Smith, K.W. & Ribi, M. (2003) The status and recent breeding performance of the critically endangered Northern Bald Ibis Geronticus eremita population on the Atlantic coast of Morocco. Ibis, 145, 419–431.

Bowden, C.G.R., Hamoud, A., Jbour, S., Fritz, J., Peske, L., Riedler, B., et al. (2012) Attempted supplementation of the relict wild Eastern population of Northern Bald Ibis in Syria with Turkish semi-wild juveniles. IUCN Reintroduction Specialists Group Case Studies Part III, 130–134.

Bowden, C.G.R., Smith, K.W., Bekkay, M. El, Oubrou, W., Aghnaj, A. & Jimenez-Armesto, M. (2008) Contribution of research to conservation action for the Northern Bald Ibis Geronticus eremita in Morocco. Bird Conservation International, 18, 74–90.

Corlett, R.T. (2016) Restoration, Reintroduction, and Rewilding in a Changing World. Trends in Ecology and Evolution.

Destro, G.F.G., De Marco, P. & Terribile, L.C. (2018) Threats for bird population restoration: A systematic review. Perspectives in Ecology and Conservation, 16, 68–73.

Frey, H. (1992) Die Wiedereinbürgerung des Bartgeiers (Gypaetus barbatus) in den Alpen. Egretta, 35, 85–95.

Fritz, J. (2015) Reintroduction of the Northern Bald Ibis in Europe: Illegal Hunting in Italy during Autumn Migration as the Main Threat. WAZA Magazin, 31 p.

Fritz, J., Hoffmann, W. & Unsöld, M. (2016) Back into European ecosystems□: The LIFE + Northern Bald Ibis reintroduction project in Central Europe. In Report of 4th IAGNBI Meeting Seekirchen (Ed. C Böhm & C Bowden).

Fritz, J., Kramer, R., Hoffmann, W., Trobe, D. & Unsöld, M. (2017) Back into the wild: establishing a migratory Northern bald ibis Geronticus eremita population in Europe. International Zoo Yearbook, 51, 107–123.

Fritz, J. & Riedler, B. (2010) Neue Hoffnung für das Überleben einer hoch bedrohtesten Zugvogelart im Mittleren Osten: Freisetzung von Jungvögeln bei den letzten migrierenden Waldrappen in Syrien. Vogelwarte, 48, 417–418.

Fritz, J., Unsoeld, M. & Voelkl, B. (2019) Back into European Wildlife: The Reintroduction of the Northern Bald Ibis (Geronticus eremita). In Scientific Foundations of Zoos and Aquariums: Their Role in Conservation and Research (eds A. Kaufman, M. Bashaw & T. Maple), pp. 339–366. Cambridge University Press.

Fritz, J., Wirtz, S. & Unsoeld, M. (2017) Aspekte der Nahrungsökologie und Genetik des Waldrapps: Reply zu Bauer et al. (2016) Vogelneozoen in Deutschland - Revision der nationalen Statuseinstufungen. Vogelwarte, 55, 141–145.

Gray, T.N.E., Crouthers, R., Ramesh, K., Vattakaven, J., Borah, J., Pasha, M.K.S., et al. (2017) A framework for assessing readiness for tiger Panthera tigris reintroduction: a case study from eastern Cambodia. Biodiversity and Conservation, 26, 2383–2399.

Grimm, V., Augusiak, J., Focks, a., Frank, B.M., Gabsi, F., Johnston, A.S.A., et al. (2014) Towards better modelling and decision support: Documenting model development, testing, and analysis using TRACE. Ecological Modelling, 280, 129–139. Elsevier B.V.

Grimm, V., Berger, U., Bastiansen, F., Eliassen, S., Ginot, V., Giske, J., et al. (2006) A standard protocol for describing individual-based and agent-based models. Ecological Modelling, 198, 115–126.

Grimm, V., Berger, U., DeAngelis, D.L., Polhill, J.G., Giske, J. & Railsback, S.F. (2010) The ODD protocol: A review and first update. Ecological Modelling, 221, 2760–2768.

Houser, A.M., Gusset, M., Bragg, C.J., Boast, L.K. & Somers, M.J. (2011) Pre-release hunting training and post-release monitoring are key components in the rehabilitation of orphaned large felids. African Journal of Wildlife Research, 41, 11–20.

IUCN/SSC (2013a) Guidelines for Reintroductions and Other Conservation Translocations. IUCN Species Survival Commission, Gland, Switzerland.

IUCN/SSC (2013b) Guidelines for Reintroductions and Other Conservation Translocations. Version 1.0. Gland, Switzerland.

Kaplan, E.L. & Meier, P. (1958) Nonparametric Estimation from Incomplete Observations. Journal of the American Statistical Association, 53, 457–481.

Kleinbaum, D.G. & Klein, M. (2012) Survival analysis: A Self-Learning Text, 3rd edition. Springer Science+Business Media.

Lampila, S., Orell, M., Belda, E. & Koivula, K. (2006) Importance of adult survival, local recruitment and immigration in a declining boreal forest passerine, the willow tit Parus montanus. Oecologia, 148, 405–413.

Lebreton, J.D. & Clobert, J. (1991) Bird population dynamics, management, and conservation: the role of mathematical modelling. In Bird population studies (eds C.M. Perrins, J.D. Lebreton & G.J.M. Hirons), pp. 105–125. Oxford University Press, Oxford.

Mathews, F., Orros, M., McLaren, G., Gelling, M. & Foster, R. (2005) Keeping fit on the ark: Assessing the suitability of captive-bred animals for release. Biological Conservation, 121, 569–577.

Muñoz, A.-R. & Ramírez, J. (2017) Reintroduced northern bald ibises from Spain reach Morocco. Oryx, 51, 204–205.

O’Shea, T.J., Ellison, L.E. & Stanley, T.R. (2011) Adult survival and population growth rate in Colorado big brown bats (Eptesicus fuscus). Journal of Mammalogy, 92, 433–443.

Parker, K.A., Ewen, J.G., Seddon, P.J. & Armstrong, D.P. (2013) Post-release monitoring of bird translocations: Why is it important and how do we do it? Notornis, 60, 85–92.

Pereira, H.M. & Navarro, L.M. (eds) (2015) Rewilding European Landscapes. In Rewilding European Landscapes p., 1st edition. Springer International Publishing, Cham.

Pettorelli, N., Barlow, J., Stephens, P.A., Durant, S.M., Connor, B., Schulte to Bühne, H., et al. (2018) Making rewilding fit for policy. Journal of Applied Ecology, 55, 1114–1125.

Pistorius, P.A., Follestad, A. & Taylor, F.E. (2006) Declining winter survival and fitness implications associated with latitudinal distribution in Norwegian Greylag Gesse Anser anser. Ibis, 148, 114–125.

Portugal, S.J., Hubel, T.Y., Fritz, J., Heese, S., TRobe, D., Voelkl, B., et al. (2014) Upwash exploitation and downwash avoidance by flap phasing in ibis formation flight. Nature, 505, 399–402.

R Core TEam (2020) R: A language and environment for statistical computing. R Foundation for Statistical Computing, Vienna, Austria.

Robert, A., Colas, B., Guigon, I., Kerbiriou, C., Mihoub, J.B., Saint-Jalme, M. & Sarrazin, F. (2015) Defining reintroduction success using IUCN criteria for threatened species: A demographic assessment. Animal Conservation, 18, 397–406.

Schaub, M., Kania, W. & Köppen, U. (2005) Variation of primary production during winter induces synchrony in survival rates in migratory white storks Ciconia ciconia. Journal of Animal Ecology, 74, 656–666.

Schaub, M., Zink, R., Beissmann, H., Sarrazin, F. & Arlettaz, R. (2009) When to end releases in reintroduction programmes: Demographic rates and population viability analysis of bearded vultures in the Alps. Journal of Applied Ecology, 46, 92–100.

Serra, G. (2015) The Northern Bald Ibis is extinct in the Middle East - but we can’t blame it on IS. Ecologist.

Serra, G., Lindsell, J.A., Peske, L., Fritz, J., Bowden, C.G.R., Bruschini, C., et al. (2014) Accounting for the low survival of the Critically Endangered northern bald ibis Geronticus eremita on a major migratory flyway. Oryx, 49, 312–320.

Smith, K.W., Aghnaj, A., El Bekkay, M., Oubrou, W., Ribi, M., Armesto, M.J. & Bowden, C.G.R. (2008) The provision of supplementary fresh water improves the breeding success of the globally threatened Northern Bald Ibis Geronticus eremita. Ibis, 150, 728–734.

Soorae, P.S. (2018) Global Reintroduction Perspectives-2018; Case studies from around the globe. IUCN/SSC Re-introduction Specialist Group & Environment Agency-ABU DHABI, Gland, Swizerland.

Sperger, C., Heller, A., Voelkl, B. & Fritz, J. (2017) Flight Strategies of Migrating Northern Bald Ibises–Analysis of GPS Data During Human-led Migration Flights. AGIT ? Journalfür Angewandte Geoinformatik, 3, 62–72.

Therneau, T.M. (2015) A Package for Survival Analysis in R.

Voelkl, B. & Fritz, J. (2017) Relation between travel strategy and social organization of migrating birds with special consideration of formation flight in the northern bald ibis. Philosophical Transactions of the Royal Society B: Biological Sciences, 372, 20160235.

Wilensky, U. (1999) NetLogo. Center for Connected Learning and Computer-Based Modeling, Northwestern University, Evanston, IL.

Wimberger, K., Downs, C.T. & Perrin, M.R. (2009) Two Unsuccessful Reintroduction Attempts of Rock Hyraxes (Procavia capensis) into a Reserve in the KwaZulu-Natal Province, South Africa. African Journal of Wildlife Research, 39, 192–201.

Wirtz, S., Boehm, C., Fritz, J., Kotrschal, K., Veith, M. & Hochkirch, A. (2018) Optimizing the genetic management of reintroduction projects: Genetic population structure of the captive Northern Bald Ibis population Sarah Wirtz. Conservation Genetics, 19, 853–864.

Zhang, M., Huang, Y., Hong, M., Zhou, S., Huang, J., Li, D., et al. (2017) Impacts of man-made provisioned food on learned cub behaviours of giant pandas in pre-release reintroduction training. Folia Zoologica, 66, 58–66.

Zoufal, K., Fritz, J., Bichler, M., Kirbauer, M., Markut, T., Meran, I., et al. (2007) Feeding ecology of the Northern Bald Ibis in different habitat types: An experimental field study with handraised individuals. In Report of the 2nd IAGNBI Meeting p.

