## Supplementary Material 1-6 for "Halfway to self-sustainability: Reintroduced migratory European Northern Bald Ibises (*Geronticus eremita*) still need management interventions for population viability"

Corinna Esterer Waldrappteam Conservation & Research, Schulgasse 28, 6162 Mutters, Austria.

Ingo Kowarik Technische Universität Berlin, Department of Ecology, Chair of Ecosystem Science and Plant Ecology, Berlin, Germany. orcid.org/0000-0002-8251-7163

Johannes Fritz (Corresponding author) Waldrappteam Conservation & Research, Schulgasse 28, 6162 Mutters, Austria. orcid.org/0000-0003-4691-2892
Also at: Department of Cognitive Biology, University of Vienna, Althanstrasse 14, 1090 Vienna, Austria

Stephanie Kramer-Schadt Department of Ecological Dynamics, Leibniz Institute for Zoo and Wildlife Research, Berlin, Germany. orcid.org/0000-0002-9269-4446
Also at: Technische Universität Berlin, Department of Ecology, Chair of Applied Animal Ecology, Berlin, Germany.

Content

### Supplementary Material 1: Study species

The Northern Bald Ibis (*Geronticus eremita,* hereafter NBI) is one of the most endangered bird species in the world: According to the IUCN Red List of Threatened Species, the NBI was critically endangered from 1994 (BirdLife International, 2017); since 2018 the species is listed as endangered due to management actions applied to the existing populations (BirdLife International, 2018).

The NBI is a migratory, mainly insectivorous, ibis species that reaches an age of up to 30 year in captivity (Böhm & Pegoraro, 2011). NBI breed in pairs that form each season and raise together up to four chicks in colonies of up to hundreds of birds (Fritz et al., 2017). Historic breeding grounds reached all around the Mediterranean, from Afrika to the Arabian Peninsula and Europe (Bowden et al., 2003; Bowden, 2015; Fritz & Unsöld, 2015; BirdLife International, 2017; Fritz et al., 2017; Böhm et al., 2020). A recent genetic study indicates a formerly contiguous population, only the extinction of the European population in the 17^th^ century separated a western and eastern population. The study found no evidence of differentiation of two evolutionary significant units (Wirtz et al., 2018). An International Single Species Action Plan (ISSAP) was established to formulate conservation goals for the NBI in 2015 after years of consultation (Bowden, 2015). The ISSAP aims to increase the population size and breeding range of the species by means of improved survival and reproduction and the establishment of new colonies.

In autumn, all individuals migrate to the wintering grounds and they return to the same breeding area the next spring. Juveniles usually stay in the wintering grounds until they are adults (Fritz et al., 2019). Worldwide, only one wild, mainly sedentary population is remaining, with two breeding sites on the Atlantic coast in Morocco (Bowden et al., 2008; Schenker et al., 2020). A relict population in the Middle East, which bred in Syria and wintered in Ethiopia, went extinct in 2013 (Serra, 2015). In Turkey, a managed, sedentary population is remaining (Yeníyurt et al., 2017). Until the 17th century, NBI bred in Europe during summer, especially around the Alps or Andalusia (Schenker, 1977; Unsöld & Fritz, 2011; Bowden, 2015; Fritz et al., 2017). The European NBI population was migratory but the historical winter areas are unknown (Fritz et al., 2017; Schenker et al., 2020). The species went extinct in Europe due to human actions, especially hunting. Eradication was probably accelerated by the little ice age and the beginning of the Thirty Years' War (Schenker, 1977; Fritz et al., 2017). Currently, there are two on-going reintroduction projects in Europe. *Proyecto Eremita* reintroduce a sedentary NBI population in Andalusia (Spain; Bowden, 2015; López & Quevedo, 2016; Böhm et al., 2020), while the so called *Waldrappteam* establish a migratory population, with breeding sites North of the Alps and a wintering ground in southern Tuscany (Fritz et al., 2019).

A group of scientists headed by J. Fritz founded the *Waldrappteam* in 2002. In the course of a feasibility study along the IUCN reintroduction guidelines (2002-2013) and during a subsequent LIFE+ biodiversity project (2014-2019; LIFE+12-BIO_AT_000143) the team released a total of 223 juveniles and reduced non-natural threats such as poaching and electrocution (Fritz et al., 2019). End of 2019, the Waldrappteam NBI population consisted of 143 individuals, which belong to four breeding colonies: Two well established colonies in Burghausen (Bavaria, Germany) and Kuchl (Salzburg, Austria) with 37 fledglings in 2019 and two colonies which consisted only of released and not yet adult birds in Überlingen (Baden-Württemberg, Germany) and Rosegg (Kärnten, Austria). All individuals are migratory, with a migration tradition to the common wintering ground Laguna di Orbetello in the Tuscany, Italy (FIG. 1 of the main text) (Fritz, Kramer, et al., 2017; Fritz et al., 2019).

### Supplementary Fig. 1: Timelines of 250 Northern Bald Ibis (NBI) from 2008 to 2019

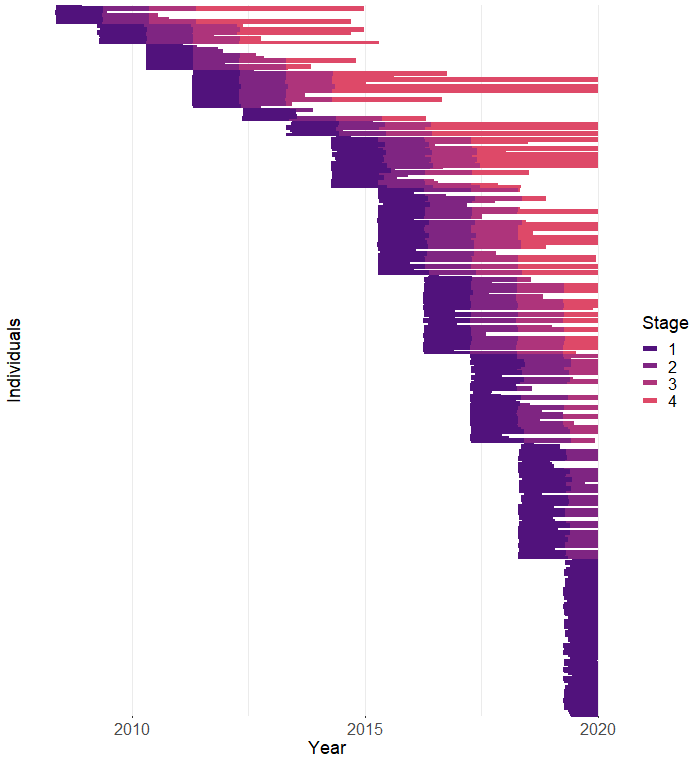

Supplementary Fig. 1 Graphical visualization of the lifetime of 384 NBI. The study period is represented on the x-axis and ranges from 2008 to 2019. Each line corresponds to one individual from its year of hatching to its year of death. The plot shows all females (N = 184), all males (N = 195) and individuals of unknown sex (N = 5) in 4 different stages (according to Figure 2) during their lifetime. Stage 1 (dark purple) corresponds to juveniles with their first migration to the wintering grounds. Stage 2 (plum) corresponds to 1 year old NBI that stay in the wintering grounds. Stage 3 (mallow) are 2 years old NBI with their first independent migration back to the breeding area. Stage 4 (coral) corresponds to reproductive adults. At the end of the data collection period the population consisted of 143 alive individuals (lines until the right border of the figure; 74 f, 69 m). 71 individuals reached only stage 1 (37 f, 34 m), 28 individuals reached only stage 2 (11 f, 17 m), 13 individuals reached only stage 3 (8 f, 5 m), 31 individuals reached stage 4 (18 f, 13 m). The legend on the right side of the plot shows the colour code for the stages considered.

### Supplementary Fig. 2: Flow chart of the study system

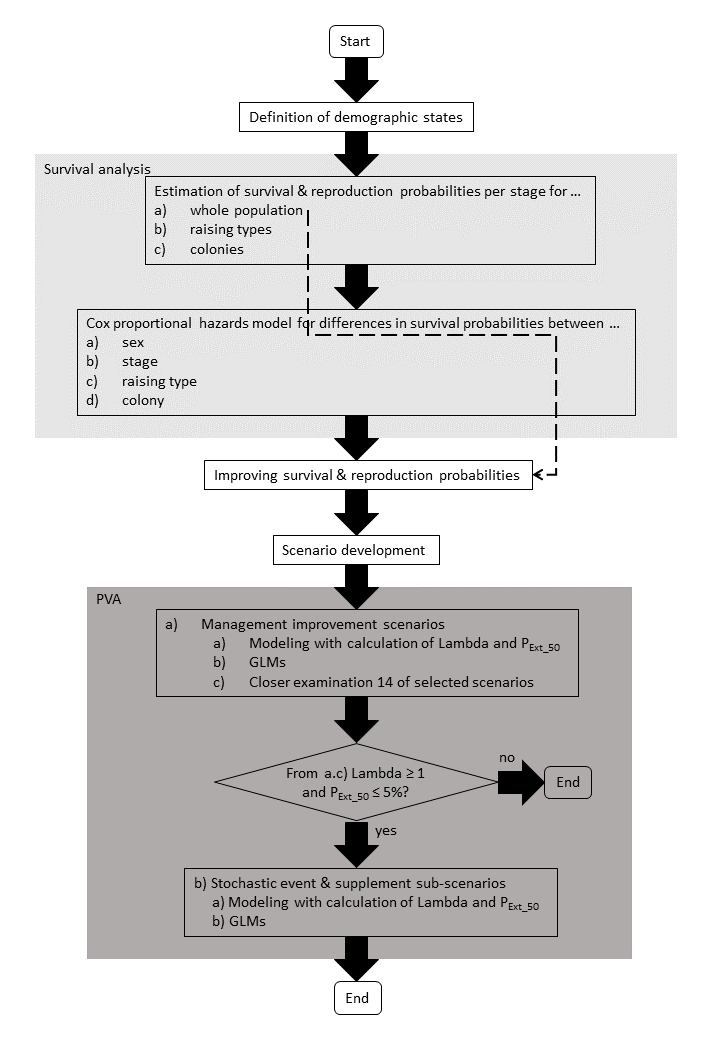

Supplementary Fig. 2 Flow chart of the study with an overview of the individual steps. Further details on the individual steps can be found in the main text. In light grey: steps of the survival analysis. In dark grey: steps of the population viability analysis (PVA). Dotted dark grey line: The estimated values for survival and reproduction are used for the steps after the survival analysis. P_EXT_50_: extinction probability after 50 years.

### Supplementary Material 2: Calculations of the different reproductive rates

#### 2.1 Calculations

##### 2.1.1 RR_Baseline_ and improvements

Here we counted only the female juveniles (raising type BP (biological parent raised)), that were born by the potential mothers (raising type BP or FP (foster parent raised)) per year and who were part of the migrating population and were not given to zoos or similar places:

$${RR}_{Baseline}=\frac{no. of female BP fledglings p.a. + \frac{1}{2} of BP fledglings of unknown sex p.a.}{no. of potential BP and FP mothers p.a.}$$

(1)

That means, also non-breeding adult females that stayed at the wintering grounds count in this analysis, hence this measure also includes the probability to breed. Additionally, we included one half of the juveniles of unknown sex. This would correspond to a scenario where any help from the Waldrappteam ended immediately. In addition, we calculated the standard deviation (SD) across years. For further scenarios regarding improvement in management e. g. in their breeding area we increased the baseline reproductive rate by 10%, 25% and 100%.

##### 2.1.2 RR_Status quo_

For this option we included the female juveniles that were born by the temporarily added females, but we did not include these females as potential mothers. This option includes all parent raised chicks per year that were part of the population and corresponds to the status quo of the population with active reproduction management:

$${RR}_{Status Quo}=\frac{\begin{aligned} total no of female BP fledglings p.a.(from potential BP and FP mothers \\ + chicks of added females)+ \frac{1}{2} of BP fledglings of unknown sex p.a. \end{aligned}}{no. of potential BP and FP mothers p.a.}$$

(2)

##### 2.1.3 RR_All chicks_

For this option we summed up the chicks of RR_Status quo_ and the female foster parent raised chicks (raising type FP) per year. Again, we divided this number of chicks by the potential mothers:

$${RR}_{All Chicks}=\frac{\begin{aligned} total no of female BP fledglings p.a. + no. of female FP fledglings p.a. \\ + \frac{1}{2} of BP fledglings of unknown sex p.a. \end{aligned}}{no. of potential BP and FP mothers p.a.}$$

(3)

##### 2.1.4 Reproductive rate per raising type

Here we counted the number of female juveniles (raising type BP), that were born by the potential mothers (raising type BP or FP) per year and per raising type of the mother and that were part of the population and were not given to zoos or similar places, like the supplementary females. The calculation was the same than for 1.1.1, but we distinguished the numbers of the chicks and the potential mothers by the raising type of the potential mothers.

##### 2.1.5 Reproductive rate per colony

Here we counted the number of female juveniles (raising type BP), that were born by the potential mothers (raising type BP or FP) per year and per colony and who were part of the population and were not given to zoos or similar places. The calculation was the same as for 1.1.1, but we distinguished the numbers of the chicks and the potential mothers by the colony of the potential mothers.

#### 2.2 Results

For RR_Baseline_ and RR_Status quo_ we did not start before 2012 with the calculations of RR, because there were no potential mothers before 2011, and in 2011 there was only one potential mother. For RR_All chicks_ we did not include the years 2011 to 2013, as there was only one potential mother in 2011 and no FP chicks were added in 2012 and 2013.

##### 2.2.1 RR_Baseline_ and improvements

We calculated a baseline reproductive rate of 0.53 (± 0.17) for a total of 37.5 female chicks over 8 years (Supplementary Fig. 3; Supplementary Table 1). Then we increased this value by 10%, 25% and 100% (Table 1 in the main text).

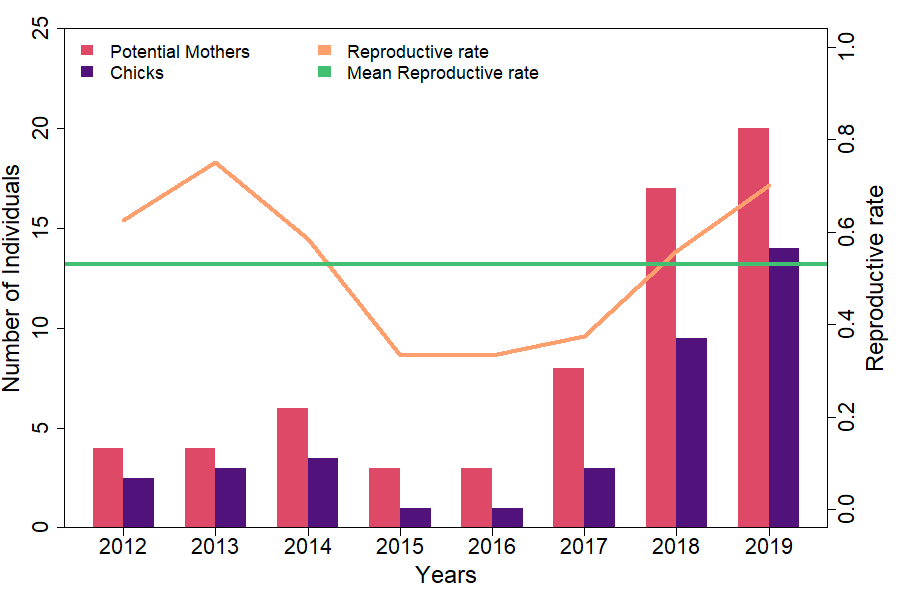

Supplementary Fig. 3 Number of potential mothers, female chicks and reproductive rate per year from 2012 to 2019 for RR_Baseline_. In addition, the mean reproductive rate across years is plotted.

##### 2.2.2 RR_Status quo_

The calculated reproductive rate for this option was 1.41 (± 0.81) for a total of 75.5 chicks over 8 years (Supplementary Fig. 4; Supplementary Table 1).

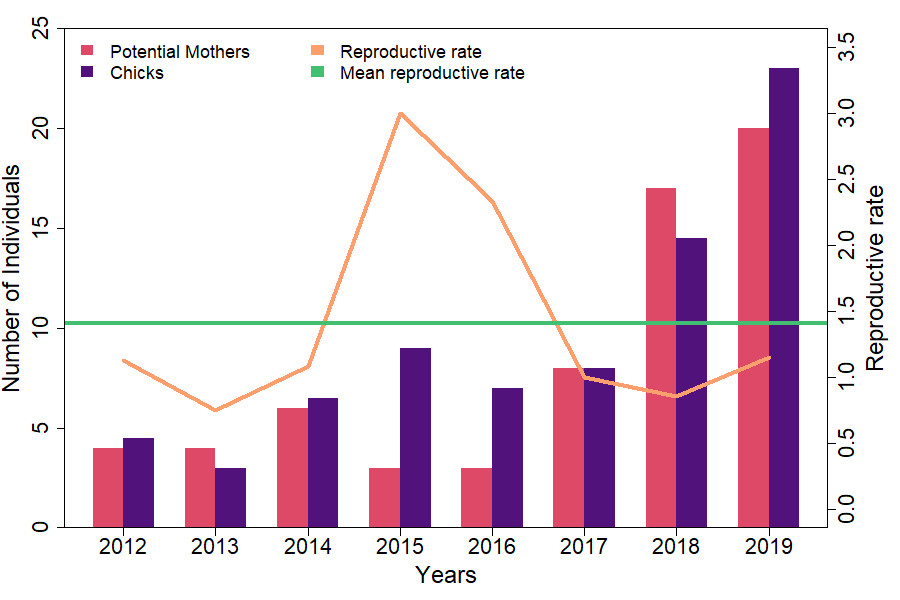

Supplementary Fig. 4 Number of potential mothers, female chicks and reproductive rate per year from 2012 to 2019 for RR_Status quo_. In addition, the mean reproductive rate across years is plotted.

##### 2.2.3 RR_All chicks_

The highest value for the reproductive rate was 3.97 (± 2.66) for a total of 151 chicks over 6 years (Supplementary Fig. 5; Supplementary Table 1).

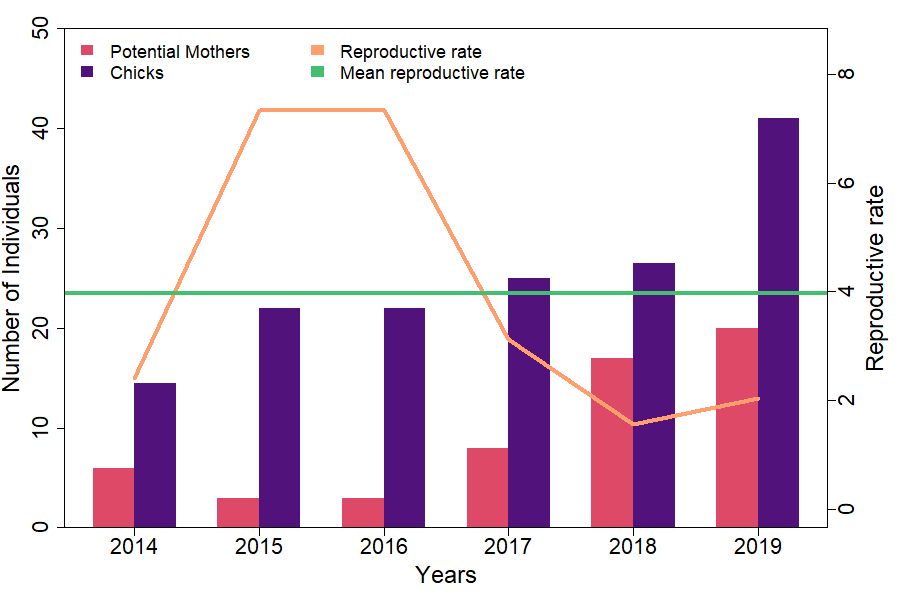

Supplementary Fig. 5 Number of potential mothers, female chicks and reproductive rate per year from 2014 to 2019 for RR_All chicks_. In addition, the mean reproductive rate across years is plotted.

The explained values for RR_Baseline_, RR_Status quo_ and RR_All chicks_ were used for the simulations with NetLogo and the BehaviorSpace-Tool (Table 1 in the main text).

Supplementary Table 1 Counted female chicks of biological parents (BP) and potential mothers (raising types BP and FP (foster parent raised)) and the calculated reproductive rate per year for RR_Baseline_, RR_Status quo_ and RR_All chicks_. In the number of chicks per year, half of the chicks of unknown sex were integrated, i. e. we used the sex-ratio of 1:1 to assign the chicks either to female or male. Thus, in some years this resulted in 0.5 chicks. For RR_Status quo_ we included the chicks of BP females in the population and the temporarily added females as considered female chicks. For RR_All chicks_ we included the chicks of BP females in the population and the FP raised chicks per year as considered female chicks. The mean annual RR_Baseline_ is 0.53 ± 0.17 SD across the years 2012-2019. The mean annual RR_Status quo_ is 1.41 ± 0.81 across the years 2012-2019. The mean annual RR_All chicks_ is 3.97 ± 2.66 across the years 2014-2019.

| Category/Year | 2012 | 2013 | 2014 | 2015 | 2016 | 2017 | 2018 | 2019 |
| --- | --- | --- | --- | --- | --- | --- | --- | --- |
| Potential mothers (BP, FP) | 4 | 4 | 6 | 3 | 3 | 8 | 17 | 20 |
| Considered female chicks for RR_Baseline_ (BP) | 2.50 | 3.00 | 3.50 | 1.00 | 1.00 | 3.00 | 9.50 | 14.00 |
| Resulting RR_Baseline_ | 0.63 | 0.75 | 0.58 | 0.33 | 0.33 | 0.38 | 0.56 | 0.70 |
| Considered female chicks for RR_Status quo_ (BP) | 4.50 | 3.00 | 6.50 | 9.00 | 7.00 | 8.00 | 14.50 | 23.00 |
| Resulting RR_Status quo_ | 1.13 | 0.75 | 1.08 | 3.00 | 2.33 | 1.00 | 0.85 | 1.15 |
| Considered female chicks for RR_All chicks_  (BP, FP) |  |  | 14.50 | 22.00 | 22.00 | 25.00 | 26.50 | 41.00 |
| Resulting RR_All chicks_ |  |  | 2.42 | 7.33 | 7.33 | 3.13 | 1.56 | 2.05 |

##### 2.2.4 Reproductive rate per raising type

We calculated a reproductive rate for FP NBI (RR_FP_) of 0.56 (± 0.14) for a total of 24 potential mothers and 30 female fledglings over 8 years (Supplementary Table 2). Additionally, we calculated a reproductive rate for BP NBI (RR_BP_) of 0.34 (± 0.31) for a total of 7 potential mothers and 5.5 female fledglings over 3 years (Supplementary Table 2).

Supplementary Table 2 Counted female chicks of biological parents (BP) and potential mothers and the calculated reproductive rate per year for biological parent raised NBI (BP) and foster parent raised NBI (FP). In the number of female chicks per year, half of the chicks of unknown sex were integrated, i. e. we used the sex-ratio of 1:1 to assign the chicks either to female or male, Thus in some years this resulted in 0.5 chicks. The mean annual reproductive rate of FP NBI is 0.56 ± 0.14 SD across the years 2012-2019 with a total of 24 potential mothers and 30 female fledglings. The mean annual reproductive rate of BP NBI is 0.34 ± 0.31 SD across the years 2017-2019 with a total of 7 potential mothers and 5.5 female fledglings.

| Category/Year | 2012 | 2013 | 2014 | 2015 | 2016 | 2017 | 2018 | 2019 |
| --- | --- | --- | --- | --- | --- | --- | --- | --- |
| Potential mothers FP | 4 | 4 | 6 | 3 | 2 | 4 | 11 | 15 |
| Female chicks (only BP) FP mothers | 2.00 | 2.00 | 3.00 | 1.00 | 1.00 | 3.00 | 7.00 | 11.00 |
| Resulting RR_FP_ | 0.50 | 0.50 | 0.50 | 0.33 | 0.50 | 0.75 | 0.64 | 0.73 |
| Potential mothers BP |  |  |  |  |  | 4 | 6 | 5 |
| Female chicks (only BP) BP mothers |  |  |  |  |  | 0 | 2.50 | 3.00 |
| Resulting RR_BP_ |  |  |  |  |  | 0 | 0.42 | 0.60 |

##### 2.2.5 Reproductive rate per colony

We calculated a reproductive rate for the colony in Burghausen (RR_B_) of 0.59 (± 0.40) for a total of 13 potential mothers and 22.5 female fledglings over 8 years (Supplementary Table 3). Additionally, we calculated a reproductive rate for the colony in Kuchl (RR_K_) of 0.42 (± 0.26) for a total of 12 potential mothers and 15 female fledglings over 6 years (Supplementary Table 3).

Supplementary Table 3 Counted female chicks of biological parents (BP) and potential mothers and the calculated reproductive rate per year for the NBI colonies in Burghausen (B) and Kuchl (K). In the number of female chicks per year, half of the chicks of unknown sex were integrated, i. e. we used the sex-ratio of 1:1 to assign the chicks either to female or male, Thus in some years this resulted in 0.5 chicks. The mean annual reproductive rate of the colony in Burghausen is 0.59 ± 0.40 SD across the years 2012-2019 with a total of 13 potential mothers and 22.5 female fledglings. The mean annual reproductive rate of the colony in Kuchl is 0.42 ± 0.26 SD across the years 2014-2019 with a total of 12 potential mothers and 15 female fledglings.

| Category/Year | 2012 | 2013 | 2014 | 2015 | 2016 | 2017 | 2018 | 2019 |
| --- | --- | --- | --- | --- | --- | --- | --- | --- |
| Potential mothers B | 4 | 4 | 3 | 0 | 1 | 5 | 6 | 6 |
| Female chicks (only BP) B | 2.50 | 3.00 | 2.50 | 0 | 0 | 3.00 | 4.50 | 7.00 |
| Resulting RR_B_ | 0.63 | 0.75 | 0.83 | 0 | 0 | 0.60 | 0.75 | 1.17 |
| Potential mothers K |  |  | 3 | 3 | 2 | 3 | 9 | 9 |
| Female chicks (only BP) K |  |  | 1.00 | 1.00 | 1.00 | 0 | 5.00 | 7.00 |
| Resulting RR_K_ |  |  | 0.33 | 0.33 | 0.50 | 0 | 0.56 | 0.78 |

### Supplementary Material 3: TRACE document according to Grimm et al. 2014.

Can be found in an extra document.

### Supplementary Material 4: Equations for the number of scenarios in NetLogo

#### 4.1 Management improvement scenarios

| No. Scenarios = 3 possible values for s1 * 3 possible values for s2 * 3 possible   values for s3 * 3 possible values for s4 * 4 possible values for   RR + “Status quo”-Scenario + “All chicks”-Scenario = 326 | (4), |
| --- | --- |

where s1 = juveniles with their first migration to the wintering grounds, s2 = 1-year-old NBI that stay in the wintering grounds, s3 = 2-year-old birds with the first independent migration back to the breeding area, s4 = reproductive adults, RR = RR_Baseline_ and the improvements.

#### 4.2 Stochastic event and juvenile supplement sub-scenarios

| No. cases = 1a * 4 freq * 5 severity +   10 sc * 4 freq * 5 severity * 2 supplements * 2 time = 820 | (5), |
| --- | --- |

where a = all chicks scenario, freq = possible values for stochastic event frequency, severity = possible values for stochastic event severity, sc = Baseline scenario and 9 out of 14 scenarios of special interest where lambda >1 and extinction probability ≤ 5%, supplements = possible values for number of supplements, time = possible durations of supplementing individuals. All possible combinations were tested.

### Supplementary Material 5: Description of the generalized linear models (GLMs)

#### 5.1 Management improvement scenarios

We tested how lambda was influenced by the survival of juvenile NBI (s1), 1-year old NBI (s2), 2-years old NBI (s3), adult NBI (s4) and the reproductive rate (RR). In this analysis, we have taken into account the mean values of 100 runs per scenario for all 326 scenarios (N = 326). The model was formulated as generalized linear model with gamma error distribution and identity link function in R-Studio. Additionally, we formulated a general additive linear model. The formulas as R code were as follows, where k is the number of smoothing splines:

| *glm* | *Lambda ~ s1 + s2 + s3 + s4 + RR* | (6) |
| --- | --- | --- |
| *gam* | *Lambda* ~ s(s1, k = 3) + s(s2, k = 3) + s(s3, k = 3) +   s(s4, k = 3) + s(RR, k = 3) | (7) |

All explanatory variables had a significant effect on lambda, but s4 had the strongest effect (Supplementary Table 4; Supplementary Table 5). The effect plots of the glm and perspective plots of the gam of the explanatory variables confirm this result (Supplementary Fig. 6; Supplementary Fig. 7). The effect plots were made with the ggeffects-package in R-Studio (Lüdecke, 2018). R^2^ is 0.97 and the effect size for a significance level of 0.05, a N of 326 and a power of 0.8 is 0.20. To calculate the effect size, we used the pwr-package in R-Studio (Champely, 2020).

Supplementary Table 4 Summary of the results of the glm-model and the effects of the explanatory variables s1 (juvenile survival), s2 (survival 1-year old NBI), s3 (survival 2-years old NBI), s4 (adult survival) and RR (reproductive rate) on lambda. t value= value of the test statistic.

| Explanatory variable | Estimate (± SE) | t value | p value |
| --- | --- | --- | --- |
| s1 | -0.20 (± 0.01) | -15.59 | <0.0001*** |
| s2 | -0.17 (± 0.01) | -14.99 | <0.0001*** |
| s3 | -0.18 (± 0.01) | -15.06 | <0.0001*** |
| s4 | -0.52 (± 0.01) | -51.15 | <0.0001*** |
| RR | 0.12 (± 0.00) | 49.43 | <0.0001*** |

Supplementary Table 5 Summary of the results of the anova of the glm-model and the effects of the explanatory variables s1 (juvenile survival), s2 (survival 1-year old NBI), s3 (survival 2-years old NBI), s4 (adult survival) and RR (reproductive rate) on lambda- Df = degrees of freedom

| Explanatory variable | Deviance | Residual Deviance | Df | Residual Df | p value |
| --- | --- | --- | --- | --- | --- |
| s1 | 0.05 | 1.45 | 1 | 324 | <0.0001*** |
| s2 | 0.05 | 1.41 | 1 | 323 | <0.0001*** |
| s3 | 0.05 | 1.36 | 1 | 322 | <0.0001*** |
| s4 | 0.68 | 0.68 | 1 | 321 | <0.0001*** |
| RR | 0.58 | 0.10 | 1 | 320 | <0.0001*** |

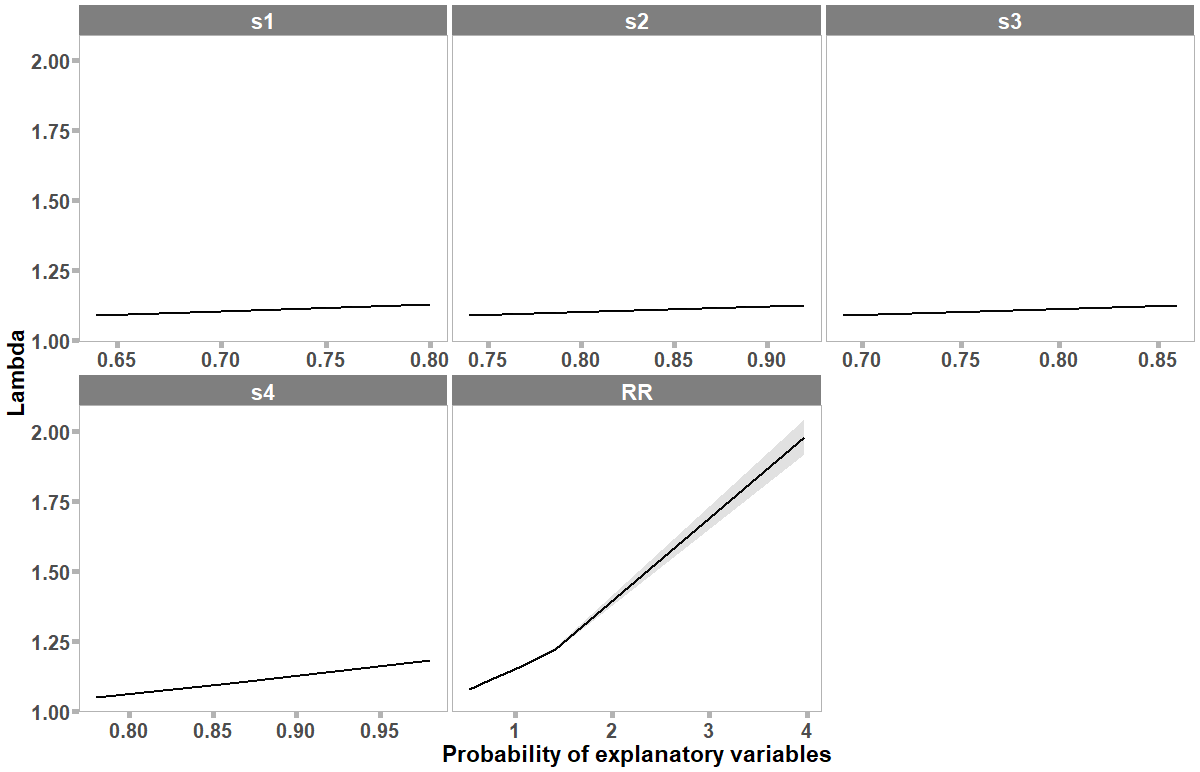

Supplementary Fig. 6 Plot of the effects of each explanatory variable of the GLM for the management improvement scenarios on the extinction probability. The explanatory variables are: s1 (juvenile survival), s2 (survival 1-year old NBI), s3 (survival 2-years old NBI), s4 (adult survival) and RR (reproductive rate). Solid line: mean effect. Blue area: confidence interval. Please consider the different x-axis and y-axis ranges.

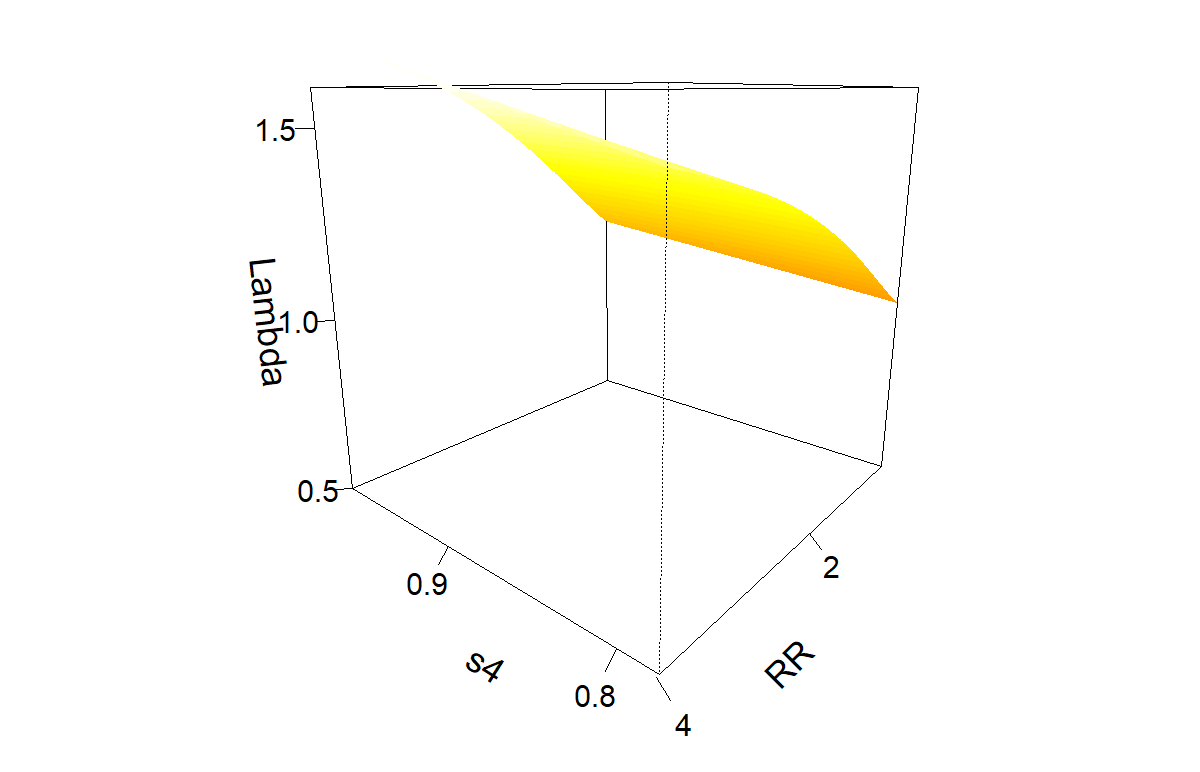

Supplementary Fig. 7 Perspective plots of the explanatory variables of a GAM for the management improvement scenarios. Plot of the effects of s4 (adult mortality) and RR (reproductive rate) on lambda. The effect plots are based on a GAM and not a GLM. However, this does not change the significance.

### Supplementary Material 6: Data and code availability

The data and the code for the analysis in R and the simulations in NetLogo can be found on: Zenodo: … The respective links will be added after the acceptance of the article.

CoMSES: … The respective links will be added after the acceptance of the article.

GitHub: … The respective links will be added after the acceptance of the article.

### Supplementary Material References

BirdLife International (2017) Geronticus eremita. *The IUCN Red List of Threatened Species 2017*.

BirdLife International (2018) Geronticus eremita. In *The IUCN Red List of Threatened Species 2018* p. .

Böhm, C., Bowden, C.G.R., Seddon, P.J., Hatipoğlu, T., Oubrou, W., El Bekkay, M., et al. (2020) The northern bald ibis Geronticus eremita : history, current status and future perspectives. *Oryx*, 1–13.

Böhm, C. & Pegoraro, K. (2011) Der Waldrapp Geronticus eremita: Ein Glatzkopf in Turbulenzen, 1st edition. VerlagsKG Wolf, Neue Brehm-Bücherei, Bd. 659, Magdeburg, Germany.

Bowden, C.G.R. (2015) International Single Species Action Plan for the Conservation of the Northern Bald Ibis (Geronticus eremita). *AEWA Technical Series No. 55*, 55.

Bowden, C.G.R., Aghnaj, A., Smith, K.W. & Ribi, M. (2003) The status and recent breeding performance of the critically endangered Northern Bald Ibis Geronticus eremita population on the Atlantic coast of Morocco. *Ibis*, 145, 419–431.

Bowden, C.G.R., Smith, K.W., El Bekkay, M., Oubrou, W., Aghnaj, A. & Jimenez-Armesto, M. (2008) Contribution of research to conservation action for the Northern Bald Ibis Geronticus eremita in Morocco. *Bird Conservation International*, 18, S74–S90. Cambridge University Press.

Champely, S. (2020) pwr: Basic Functions for Power Analysis v1.3-0. Https://cran.r-project.org/package=pwr.

Fritz, J., Kramer, R., Hoffmann, W., Trobe, D. & Unsöld, M. (2017) Back into the wild: establishing a migratory Northern bald ibis Geronticus eremita population in Europe. *International Zoo Yearbook*, 51, 107–123.

Fritz, J. & Unsöld, M. (2015) Internationaler Artenschutz im Kontext der IUCN Reintroduction Guidelines: Argumente zur Wiederansiedlung des Waldrapps Geronticus Eremita in Europa. *Vogelwarte*, 53, 157–168.

Fritz, J., Unsöld, M. & Völkl, B. (2019) Back into European Wildlife: the reintroduction of the northern bald ibis (Geronticus eremita). In *Scientific Foundations of Zoos and Aquariums: their Role in Conservation and Research* (eds A.B. Kaufman, M.J. Bashaw & T.L. Maple), pp. 339–366, 1st edition. Cambridge University Press, Cambridge, UK.

López, J.M. & Quevedo, M.A. (2016) Northern Bald Ibis Reintroduction program in Andalusia. In *Proceedings of 4th International Advisory Group for the Northern Bald Ibis (IAGNBI) meeting Seekirchen, Austria* (eds C. Böhm & C.G.R. Bowden), pp. 57–67. RSPB, The Lodge, Sandy, Bedfordshire, UK and Alpenzoo Innsbruck-Tirol, Austria.

Lüdecke, D. (2018) ggeffects: Tidy Data Frames of Marginal Effects from Regression Models. *Journal of Open Source Software*, 3, 772.

Schenker, A. (1977) Das ehemalige Verbreitungsgebiet des Waldrapps Geronticus eremita in Europa. *Der Ornithologische Beobachter*, 74, 13–30.

Schenker, A., Cahenzli, F., Gutbrod, K.G., Thevenot, M. & Erhardt, A. (2020) The Northern Bald Ibis Geronticus eremita in Morocco since 1900: Analysis of ecological requirements. *Bird Conservation International*, 30, 117–138. Cambridge University Press.

Serra, G. (2015) The Northern Bald Ibis is extinct in the Middle East - but we can’t blame it on IS. *Ecologist*.

Unsöld, M. & Fritz, J. (2011) Der Waldrapp-ein Vogel zwischen Ausrottung und Wiederkehr. *Wildbiologie 2: 1-16.*, 2, 1–16.

Wirtz, S., Böhm, C., Fritz, J., Kotrschal, K., Veith, M. & Hochkirch, A. (2018) Optimizing the genetic management of reintroduction projects: genetic population structure of the captive Northern Bald Ibis population. *Conservation Genetics*, 19, 853–864.

Yeníyurt, C., Oppel, S., Ísfendíyaroğlu, S., Özkinaci, G., Erkol, I.L. & Bowden, C.G.R. (2017) Influence of feeding ecology on breeding success of a semi-wild population of the critically endangered Northern Bald Ibis Geronticus eremita in southern Turkey. *Bird Conservation International*, 27, 537–549.
