## Supplementary Material 3 TRACE for "Halfway to self-sustainability: Reintroduced migratory European Northern Bald Ibises (*Geronticus eremita*) still need management interventions for population viability"

CORINNA ESTERER Waldrappteam Conservation & Research, Schulgasse 28, 6162 Mutters, Austria.

INGO KOWARIK Technische Universität Berlin, Department of Ecology, Chair of Ecosystem Science and Plant Ecology, Berlin, Germany. orcid.org/0000-0002-8251-7163

JOHANNES FRITZ (Corresponding author) Waldrappteam Conservation & Research, Schulgasse 28, 6162 Mutters, Austria. orcid.org/0000-0003-4691-2892

Also at: Department of Cognitive Biology, University of Vienna, Althanstrasse 14, 1090 Vienna, Austria

STEPHANIE KRAMER-SCHADT Department of Ecological Dynamics, Leibniz Institute for Zoo and Wildlife Research, Berlin, Germany. orcid.org/0000-0002-9269-4446

Also at: Technische Universität Berlin, Department of Ecology, Chair of Applied Animal Ecology, Berlin, Germany.

**Supplementary Material 3: TRACE document according to Grimm et al. 2014.**

**TRACE document**

This is a TRACE document (“TRAnsparent and Comprehensive model Evaludation”) which provides supporting evidence that our model presented in:

<**UPDATED UPON ACCEPTANCE**>

was thoughtfully designed, correctly implemented, thoroughly tested, well understood, and appropriately used for its intended purpose. The rationale of this document follows:

Schmolke A, Thorbek P, DeAngelis DL, Grimm V. 2010. Ecological modelling supporting environmental decision making: a strategy for the future. *Trends in Ecology and Evolution* 25: 479-486.

and uses the updated standard terminology and document structure in:

Grimm V, Augusiak J, Focks A, Frank B, Gabsi F, Johnston ASA, Kułakowska K, Liu C, Martin BT, Meli M, Radchuk V, Schmolke A, Thorbek P, Railsback SF. 2014. Towards better modelling and decision support: documenting model development, testing, and analysis using TRACE. *Ecological Modelling 280:* 129-139.

and

Augusiak J, Van den Brink PJ, Grimm V. 2014. Merging validation and evaluation of ecological models to ‘evaludation’: a review of terminology and a practical approach. *Ecological Modelling 280:* 117-128.

### 3.1 Model description

Summary: The purpose of the model is to investigate the population viability of the reintroduced Northern Bald Ibis (NBI) population. We tested how the demographic effects of survival and reproduction probabilities for the four different stages, as analysed from field data, changed the model predictions about the population viability. Only females are considered in the model.

#### 3.1.1 Purpose

The purpose of the model is to investigate if the reintroduced NBI population is viable. We tested how different survival probabilities and reproduction probabilities changed the model predictions.

#### 3.1.2 Entities, state variables, and scales

The model entities are female individuals in four different stages (Supplementary Table 3.1). All individuals are described by constant state variables, Raising type (raising type in Stage 1), and their age (Years of life). One step in the simulation corresponds to one year. Each run takes 50 years, and we did 100 repetitions per run. Space is not considered.

#### 3.1.3 Process overview and scheduling

At each time step, a year, the entities, female NBI, go through the processes in the following order: *breeding*, *death* and *aging*.
*Breeding*: Only the adults, NBI in stage 4, can reproduce at the beginning of each time step and produce female chicks. For the simulation of the baseline scenario, we included the standard deviation across years, so the reproductive rate varies in each time step, i. e. year, adding environmental variability such as good or bad years to the simulations.

*Death*: For every stage there is a certain mortality probability, the opposite of our calculated survival probabilities and their improvements (Supplementary Table 3.2). We included the standard deviation only for the simulation of the baseline scenario, described under 3.1.7.2.2.

*Aging*: The state variable age will be updated during the *aging* process synchronously for all individuals at the end of each time step. All individuals who reach the age of 25 years will be removed (‘die’) as this is the maximum lifespan of an NBI (Bowden, 2015).

Supplementary Table 3.1 Entities, their corresponding state variables and possible status or units. In brackets: Notation in the NetLogo Code.

| Entity (code notation) | State variable (code notation) | Status/Units |
| --- | --- | --- |
| Stage 1 Juveniles: Fledgling and first-time migrator, either with foster parents or biological parents  For BP: first-time migration back to breeding area alone in spring  (Number_Juveniles) | Age (age)  Raising type (raising) | Numeric (years)  FP (foster parent raised),  BP (biological parent raised) |
| Stage 2 1-Year-Old : for BP: experienced migrator; for FP: in wintering grounds  (Number_Subadults_Age1) | Age (age)  Raising type (raising) | Numeric (years)  FP (foster parent raised),  BP (biological parent raised) |
| Stage 3 2-Year-Old: for BP: experienced migrator; for FP: in wintering grounds and first-time migration back to breeding area alone  (Number_Subadults_Age2) | Age (age)  Raising type (raising) | Numeric (years)  FP (foster parent raised),  BP (biological parent raised) |
| Stage 4 Adults: reproductive age class and experienced migrator  (Number_Adults) | Age (age)  Raising type (raising) | Numeric (years)  FP (foster parent raised),  BP (biological parent raised) |

#### 3.1.4 Design concepts

*Basic principles.* The model is based on the demographic effects of mortality and reproduction probabilities for different stages as analysed from field data.

*Emergence.* The population trajectories emerge from the underlying processes of mortality and reproduction and may vary because of different survival and reproduction probabilities. Even with the same probabilities different results are possible due to environmental stochasticity.

*Sensing.* NA

*Interaction.* There are no interactions between agents. Only females are considered

*Stochasticity.* We have stochasticity in the birth and death processes. We took a random number for each individual and compared it to the number of the death probability for the respective stage of the individual. If the random was number smaller, the agent died. And we took a random number for each individual in stage 4 and compared it with the value for the reproduction. If the random number was smaller, the agent reproduced. And only for the simulation of the baseline scenario did we additionally implement the standard deviation for the survival and reproduction probabilities. Thus, the survival and reproduction probabilities changed at each time step and were within one standard deviation. Then we proceeded as described above.

*Collectives.* The individuals are assigned to 2 different raising types and 3 different colonies (Supplementary Table 3.1). In this model, for the simulation for H1, the different groups have no effects on survival and reproduction. But for H2 and H3 we calculated different survival and reproduction probabilities depending on the raising type (H2) or the colony (H3), as explained in the main text.

*Observation.* Number of individuals, individuals per raising type, individuals per stage. All numbers were gathered at each time step. Please note that we only modelled the female part of the population.

#### 3.1.5 Initialization

The values for the female start population at time t = 0 are 37 juveniles, 11 One-Year-Old NBI, 8 Two-Year-Old NBI and 18 adults. The values for the start population were taken from the field data from the Waldrappteam.

Supplementary Table 3.2 Parameters used for the individual-based model in NetLogo. Parameters are described with their definition, baseline value in the simulation (baseline scenario), other possible values (in different combinations in the other scenarios) and unit. In brackets in the column default value: standard deviation (SD).

| Name | Definition | Baseline value (±SD) | Other possible values | Unit |
| --- | --- | --- | --- | --- |
| Number_Juveniles | Number of female individuals in Stage 1 | 37 | - | Number |
| Number_Subadults_Age1 | Number of female individuals in Stage 2 | 11 | - | Number |
| Number_Subadults_Age2 | Number of female individuals in Stage 3 | 8 | - | Number |
| Number_Adults | Number of female individuals in Stage 4 | 18 | - | Number |
| Mortality_Juveniles | Mortality probability of individuals in Stage 1 (s1) | 0.36 (±0.36) | 0.20, 0.30 |  |
| Mortality_Subadults_Age1 | Mortality probability of individuals in Stage 2 (s2) | 0.26 (±0.35) | 0.08, 0.19 |  |
| Mortality_Subadults_Age2 | Mortality probability of individuals in Stage 3 (s3) | 0.31 (±0.35) | 0.14, 0.24 |  |
| Mortality_Adults | Mortality probability of individuals in Stage 4 (s4) | 0.22 (±0.14) | 0.02, 0.14 |  |
| Repro_Rate | Probability to hatch a chick | 0.53(±0.17) | 0.58, 0.66, 1.06, 1.41, 3.97 |  |

#### 3.1.6 Input data

The model does not use input data.

#### 3.1.7 Sub models

There are three main sub models (*breeding, death, aging*) and for the baseline scenario there are six submodels (*breeding_sd, death_juveniles, death_one_year_olds, death_two_year_olds, death_adults, aging*).

##### 3.1.7.1 Baseline values and Improvements

For these scenarios, which were simulated with the baseline values and the improved values for reproduction and survival, we only simulated the mean reproductive rate and survival values without including the standard deviation.

###### 3.1.7.1.1 Breeding

How many chicks a female hatches is defined through the reproductive rate, i. e. the number of fledged chicks per female. As potential mothers only individuals in Stage 4 count, because only these reached sexual maturity.

Is the reproductive rate < 1, a random number between 0-1 is drawn. If this value is smaller than the reproductive rate the female hatches a chick otherwise not. Is the reproductive rate between 1 and 2 a random number between 1 and 2 is drawn. Is the value smaller than the reproductive rate the female hatches 2 chicks otherwise 1. Is the reproductive rate between 2 and 3 a random number between 2 and 3 is drawn. Is the value smaller than the reproductive rate the female hatches 3 chicks otherwise 2. Is the reproductive rate between 3 and 4 a random number between 3 and 4 is drawn. Is the value smaller than the reproductive rate the female hatches 4 chicks otherwise 3.
All born chicks have the raising type “BP”.

###### 3.1.7.1.2 Death

For each stage there is a unique mortality probability (Mortality_Juveniles, Mortality_Subadults_Age1, Mortality_Subadults_Age2, Mortality_Adults). Each stage is defined through the age, and for each individual in one of the stages a random number between 0 and 1 is drawn. Is this number smaller than the respective mortality probability the individual dies.

###### 3.1.7.1.3 Aging

At the end of each time step the individuals age one year. Individuals from stage 1, 2 or 3 reach the next stage. Individuals in stage 4 remain in this stage until the end of their life, but their age status is updated. Chicks are in the first stage. Individuals older than 25 years will die.

##### 3.1.7.2 Baseline scenario

For this scenario, which was simulated only with the baseline values for reproduction and survival, we included the standard deviation. Every year a new survival and reproduction probability was set for each individual. This corresponds to demographic stochasticity.

###### 3.1.7.2.1 Breeding_sd

Only females in stage 4 were included. We bounded the reproductive rates within their standard deviations as follows: A counter variable will be set to 0. While this counter variable is 0, a random number will be drawn from a normal distribution with mean = reproductive rate ±1 SD (step_repro; Supplementary Table 3.2). Is this value <1 or >0, the counter variable will be set to 1 and the while loop ends. Then a random number between 0 and 1 will be drawn. Please note that for the baseline scenario the reproductive rates were <1 (see chapter 3.1.7.1.1). Is this number smaller than step_repro the female hatches a chick, otherwise not. The step_repro will be drawn each time step for each individual. The raising type is again set to “BP”.

###### 3.1.7.2.2 death_juveniles, death_one_year_olds, death_two_year_olds, death_adults

We bounded the mortality probabilities within their standard deviations in the following way: A counter variable will be set to 0. While this counter variable is 0, a random number will be drawn from a normal distribution with mean as given by the average mortality probability for each stage ±1 SD (step_mortality). Is this value <1 or >0 the counter variable will be set to 1 and the while loop ends. Then a random number between 1 and 0 will be drawn. Is this number smaller than the step_mortality of the stage, the individual dies. The step_mortality will be drawn each time step for each individual in the respective stage accounting for demographic stochasticity.

### 3.2 Model analysis

Summary: We analysed the model in two ways: (I) management improvement scenarios and (II) stochastic event and juvenile supplement sub-scenarios.
For the management improvement scenarios, we tested the different combinations of the parameters for survival (s1-s4) and reproductive rate (RR; calculations are described in the main text).

#### 3.2.1 Management improvement scenarios

The first model analysis is the management improvement scenarios. Here we performed local and global sensitivity analyses. At first, we tested different sets, the scenarios, of the parameters for survival (s1-s4) and reproductive rate (RR) (calculations are described in the main text and Supplementary Material 2). Here either one of the parameters (local sensitivity analysis) or more were changed (global sensitivity analysis). For both ways we analysed the population trajectories on the basis of the number of individuals per scenario and per time step (year). Furthermore, we calculated the extinction probability as the number of runs of 100 repetitions per scenario where the population died out (0 individuals), and lambda, the intrinsic growth rate of the population. Scenarios where lambda > 1, which means population growth, and extinction probability ≤ 5% were chosen for more detailed analyses. We used a 5% limit as this is a commonly used limit for extinction probability of the MVP (Flather et al., 2011). The distribution of the input parameters of mortality and reproduction probabilities in scenarios where lambda > 1 and extinction probability ≤ 5% was analysed. In addition, we set up a generalized linear model (glm) to rank the effects of survival and reproduction probabilities on lambda (see also Supplementary Material 5). Beside this we chose scenarios of special interest for closer examination. These are: baseline, juvenile survival (s1) improved by 10% and 25%; adult survival (s4) improved by 10% and 25%; reproductive rate (RR) improved by 10%, 25%, 100%; 10% and 25% improved survival for all stages, 10% and 25% improved survival for all stages and for reproductive rate, “status quo” and “all chicks”.

#### 3.2.2 Stochastic event and juvenile supplement sub-scenarios

The second analysis dealt with the topics stochastic events and supplementation of FP juveniles. Stochastic events can be e. g. droughts or storms. We chose from the scenarios of special interest, the scenarios where lambda >1 and extinction probability ≤ 5%. For the supplements we assumed 15 or 30 juveniles were added per year for the duration of 4 or 7 years. The stochastic events were modelled with frequencies between 5 and 20% (5, 10, 15, 20). This corresponds to a mean frequency of every 20 years to every 5 years, respectively. The severity of the stochastic events was assumed to be between 5 to 25% (5, 10, 15, 20, 25) additional mortality per stochastic event. This resulted in 80 combinations of these values, the sub-scenarios, per scenario. We calculated lambda, the intrinsic growth rate of the population, and extinction probability, number of runs of 100 repetitions per case where the population died out (0 individuals) and analysed the distribution between the mortality rates per stage and the reproductive rate. Additionally, we set up a GLMs (see main text). lambda was the response variable, and the predictor variables were formed by the survival probabilities per stage (s1-s4), the reproductive rate (RR) and the stochastic event frequency and severity. Besides, we looked how often each combination of stochastic event frequency (5-20%) and severity (5-25%) and each combination of numbers of supplements (15 or 30) and time (4 or 7 years) were used. This is also a global sensitivity analysis where several parameters were changed simultaneously.

### 3.3 References

Bowden, C.G.R. (2015) International Single Species Action Plan for the Conservation of the Northern Bald Ibis (Geronticus eremita). *AEWA Technical Series No. 55*, 55.

Flather, C.H., Hayward, G.D., Beissinger, S.R. & Stephens, P.A. (2011) Minimum viable populations : is there a ‘ magic number ’ for conservation practitioners ? *Trends in Ecology and Evolution*, 26, 307–316. Elsevier Ltd.
